# Estradiol Influences Adenosinergic Signaling and NREM Sleep Need in Adult Female Rats

**DOI:** 10.1101/2021.05.27.445868

**Authors:** Philip C. Smith, Derrick J. Phillips, Ana Pocivavsek, Carissa A. Byrd, Shaun S. Viechweg, Brian Hampton, Jessica A. Mong

## Abstract

Studies report estradiol (E2) suppresses sleep in females; however, the mechanisms of E2 action remain largely undetermined. Our previous findings suggest that the median preoptic nucleus (MnPO) is a key nexus for E2 action on sleep. Here, using behavioral, neurochemical and pharmacological approaches, we investigated whether E2 influenced the sleep homeostat as well as adenosinergic signaling in the MnPO of adult female rats. During the Light Phase, where rats accumulate the majority of sleep, E2 markedly reduced NREM-SWA (a measure of the homeostatic sleep need). Following 6-hours of sleep deprivation, levels of NREM-SWA were significantly increased compared to baseline sleep. However, the NREM-SWA levels were not different between E2 and control treatment despite a significant increase in wake at the expense of NREM sleep. Analysis of NREM-SWA differences between baseline and recovery sleep following sleep deprivation demonstrated that E2 induced a 2-fold increase in delta power compared to controls suggesting that E2 significantly expanded the dynamic range for the sleep homeostat. Correlated with E2-induced changes in physiological markers of homeostatic sleep was a marked increase in extracellular adenosine (a molecular marker of homeostatic sleep need) during unrestricted and recovery sleep following a 6-hour deprivation. Additionally, E2 blocked the ability of an adenosine A2A receptor agonist (CGS-21680) to increase NREM sleep compared to controls. Thus, taken together, the findings that E2 increased extracellular adenosine content, while blocking A_2A_ signaling in the MnPO suggests a potential mechanism for how estrogens impact sleep in the female brain.

**Statement of Significance:** While gonadal steroids and gender are implicated as risk factors for sleep disruptions and insomnia, the relationship between ovarian steroids and sleep is poorly understood. Understanding the mechanisms through which estradiol (E2) is working to influence sleep-wake behavior is a critical first step toward a better understanding of the role of estrogens in sleep pathologies. Using a rodent model, the current study presents novel findings suggesting that estradiol (E2) is influencing adenosinergic actions in the MnPO. The ability of E2 to attenuate the local effects of the A_2A_ receptors in the MnPO suggests that E2 modulation of A_2A_ receptor signaling may underlie estrogenic suppression of sleep behavior as well as changes in homeostatic sleep need.

## Introduction

Primary sleep disorders are among the most common medical conditions; as many as one in three individuals may have sleep problems^1–3^ with an annual economic impact of over $100 billion.^4^ Clinical data show women are 40% more likely to experience one or more symptoms of insomnia over their lifespan.^5–7^ This increased risk emerges at puberty^8, 9^ and has been associated with fluctuations in ovarian steroids, particularly estrogens, suggesting that gonadal steroids and biological sex are significant risk factors for sleep disruptions.^10^ While chronic insufficient sleep is a risk factor for a variety of psychological^11–15^, neurological, and neurodegenerative pathologies^16–23^ as well as cardiovascular and metabolic dysfunctions^24–28^, clinical studies reveal that women suffering from sleep disturbances and insufficient sleep are at greater risk compared to men for mood disorders such as depression^29^ as well as metabolic^30^ and cardiovascular dysfunction.^28, 31–33^ Thus, given the increased risks to psychological and physiological well-being, sleep disorders among women are a significant public health concern.

Despite a deepening understanding of sleep regulatory mechanisms, how estrogens influence sleep circuitry is poorly understood. In women approaching menopause, the loss of estrogens is associated with insomnia, frequent night-time awakenings, and poor sleep.^10^ In contrast, recent data from young women of reproductive age suggest that the presence of estrogens are a contributing factor to sleep disturbances.^34–38^ In order to better understand the biological basis for these apparent paradoxical effects of estrogens on sleep disturbances across a women’s lifespan, it is necessary to more fully understand the mechanisms through which estrogens are influencing the sleep-circuitry.

Historically, male rodents have served as the cornerstone for elucidating the neural circuitries governing sleep. Unfortunately, this has resulted in a significant gap in our understanding of how estrogens modulate these circuits in females. Female rodents offer the opportunity to probe the sleep circuitry in order to elucidate the mechanisms by which ovarian steroids modulate sleep. Sleep patterns in the female rat are exquisitely sensitive to natural fluctuations in ovarian steroids such as estradiol (E2).^39–41^ The highly reproducible effects of exogenously administered E2 on sleep in female rodents provide an informative bioassay to investigate how E2 may be affecting sleep mechanisms. Using adult female rats, studies consistently demonstrate that sleep time is significantly reduced when endogenous ovarian steroids or exogenous E2 are elevated in females ^40, 42^ but not males^42^. Our previous findings suggest this change in sleep may be mediated through E2 actions in the preoptic area (POA) sleep switches that include the, ventrolateral preoptic area (VLPO)^40, 43, 44^ and the median preoptic nucleus (MnPO).^45^ Moreover, these POA sleep nuclei have been implicated in sensing homeostatic sleep need.^46–49^

The sleep homeostat governs the amount of sleep needed after a given period of wake to maintain homeostasis. While the mechanism and circuitry of the sleep homeostat remains elusive, sleep intensity, as measured by EEG Slow Wave Activity (0.5-4.5Hz) during NREM sleep (referred to within as NREM-SWA) and NREM sleep duration are characteristic hallmarks of sleep need. Increases in the intensity of NREM-SWA are proportional to the amount of prior wake time, while time spent in NREM sleep will dissipate the concentration of SWA.^50–55^

The neuromodulator adenosine is highly-recognized as both a sleep promoting substance and a marker of homeostatic need.^56, 57^ Adenosine levels increase proportionally with wake time and decrease during sleep in the basal forebrain and cortex.^58–60^ Furthermore, activation of adenosine receptors, A_2A_ and A_1_, in key sleep- and wake- regulating nuclei modulate sleep need and arousal.^56, 57^ Although the exact mechanisms underlying sleep homeostasis have yet to be clearly elucidated, numerous pharmacological studies and transgenic models have indicated a role for the A_1_ receptor in SWA and sleep homeostasis.^57^ Emerging evidence from the POA also suggest that A_2A_ receptors residing in the MnPO and VLPO may also play a role in sleep homeostasis.^49, 61^

While numerous independent studies in rodents clearly demonstrate that E2 markedly increases wake and suppresses NREM sleep particularly in the dark or active phase, little attention has been paid to E2 effects on sleep homeostasis in the light-phase. Indeed, previous studies demonstrate that E2 does not significantly change the duration of light phase NREM sleep; however two studies in gonadally intact cycling females have reported a significant decrease in NREM-SWA toward the end of the light-phase on proestrus when endogenous E2 levels are high.^41, 62^ Nevertheless, it is not entirely clear if increases in E2 result in changes to homeostatic sleep pressure and/or adenosine during the light or quiescent phase. Here, we seek to investigate whether interactions between E2 and the adenosine system are involved in mediating E2 effects on sleep and homeostatic sleep pressure. Overall, the present findings suggest that (1) markers of homeostatic sleep need (NREM sleep duration and SWA) were decreased in the presence of E2; (2) despite an apparent reduction in sleep need, extracellular adenosine content in the POA was markedly increased in the presence of E2 and (3) A_2A_ receptor-induced sleep was attenuated in the presence of E2. Taken together, these data suggest that E2 may be decreasing homeostatic sleep need by disrupting adenosine signaling in the MnPO.

## Materials & Methods

### Animals

Adult female Sprague–Dawley rats (250-350g) were purchased from Charles River Laboratories (Kingston, N.Y.) and housed in the Laboratory Animal Facilities at the University of Maryland, School of Medicine under a 12 h:12 h dark: light cycle. Upon arrival, animals were acclimated to the animal facility for at 7-10 days prior to the start of the experiments. Food and water were available ad libitum.

In all experiments, zeitgeber time 0 (Zt 0) represents the beginning of lights ON and Zt 12 represents the beginning of lights OFF. All experimental procedures were run in several cohorts, with all experimental groups represented in each cohort. All procedures were performed in accordance with the National Institutes of Health guide for care and use of laboratory animals and were approved by and in accordance with the guidelines of the University of Maryland Institutional Animal Care and Use Committee.

### Surgical Procedures

Surgical procedures were performed under isoflurane anesthesia with a maintenance flow of 1-3% Isoflurane with oxygen. All animals were allowed at least 7 days to recover from the surgical procedures before the start of any experiments.

#### Ovariectomy and Transmitter Implantation

Animals were ovariectomized (OVX) and simultaneously implanted with TL11M2-F40-EET transmitters (Data Sciences International, St. Paul, Minn.) in a single surgical procedure. Briefly, a midline post-dorsal skin incision (2-2.5cm) approximately halfway between the middle of the back and the base of the tail was made followed by right and left abdominal incisions (0.5 cm) through the muscle allowing for the isolation and removal of the ovaries bilaterally. Through one of the existing abdominal incisions, a bipotential-lead transmitter (DSI Inc., St. Paul, Minn.) was implanted intraperitoneally. Before closing the muscle wall, the transmitter leads were bundled together, placed in the subcutaneous space and the muscle wall closed so that the base of the transmitter leads (ends arising from the transmitter) are anchored in place.

Next, dorsal neck incision (∼3 cm) was made through the skin along the dorsal midline from the posterior margin of the eyes to a point midway between the scapulae exposing the skull and neck muscle. The bundled electrode leads were threaded through this incision to their appropriate implant sites in the neck and cranium. For cranial implantation, two burr holes were drilled asymmetrically 2 mm anterior/1.5 mm lateral and 7 mm posterior/1.5 mm lateral relative to the bregma (Figure 1A) and stainless-steel screws (Plastics One, Roanoke, Va.) were implanted. One set of leads (for the electroencephalogram; EEG) were wrapped around the screws and secured with a dental cement cap. The second set of leads (for the electromyogram; EMG) were implanted directly in the dorsal cervical neck muscle, approximately 1.0 mm apart, and sutured in place.

**Figure 1.**
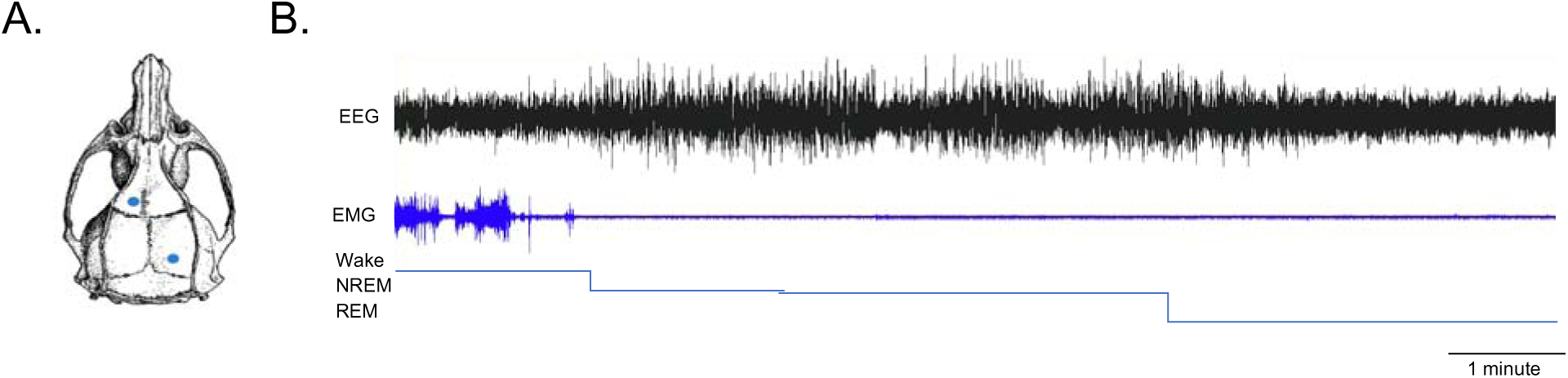
Vigilant State Scoring. (A) Configuration of recording electrode placement. Two burr holes were drilled asymmetrically and stainless-steel screws were implanted at 2 mm anterior/1.5 mm lateral and 7 mm posterior/1.5 mm lateral to Bregma. The electroencephalogram (EEG) leads were wrapped around the screws and secured with dental cement. Additionally, a second set of leads were implanted directly in the dorsal cervical neck muscle approximately 1.0 mm apart and served to record nuchal electromyograms (EMG) (B) Representative continuous recording of EEG and EMG. From the acquired continuous EEG/EMG traces, vigilance states were first automatically scored as Wake, NREM Sleep, or REM Sleep using a custom program developed in house (DJP). Additionally, experimenters blinded to the treatment groups confirmed the automated scores.

#### Guide Cannula Implantation

When necessary, for local drug infusions, guide cannula targeting the MnPO were implanted at the time of the OVX/transmitter implantation surgery. Briefly, the guide cannula (PlasticsOne #C315G, 0.46mm outer diameter, Roanoke, Va) were stereotaxically implanted after the EEG electrode placement and prior to the placement of the dental cement cap. A single burr hole was drilled in the skull bone 0.45mm posterior and 1.0mm lateral to Bregma. The guide cannula was inserted to a depth of 6.5 mm (∼0.5mm above the MnPO) at an angle of 9 degrees. The cannula and EEG leads were secured with a dental cement cap. Finally, the cannula opening was closed with a dummy cap provided by manufacturer.

For the microdialysis of MnPO interstitial fluid, animals were OVX as described above and implanted with microdialysis guide cannula (SciPro inc. #MAB-6.14.G, 1mm outer diameter, Sanborn, N.Y.) in a single surgery. Briefly, the guide cannula were stereotaxically targeted to ∼1mm above the MnPO. A single burr hole was drilled in the skull bone 0.45mm posterior and 1.0mm lateral to Bregma. The guide cannula was inserted to a depth of 6 mm at an angle of 9 degrees, secured in place with a dental cement cap and the cannula opening closed with a dummy cap provided by manufacturer. In small cohort of animals, transmitters and associated leads were implanted with microdialysis guide cannula in a single surgery as described above.

For all experiments, cannula placements were verified at the conclusions of the sample collections. Brains were collected and submersion-fixed in a solution of 9% formalin in potassium phosphate buffered saline (kPBS; 0.5M, pH 7.4), at 4°C, followed by cryoprotection in 30% sucrose in kPBS. After cryoprotection, the brains were frozen on dry ice and stored at -80 °C until processing. Brains were sectioned in 30μ coronal sections in a cryostat, directly collected on gelatin-subbed slides, dried and processed for Neutral Red staining. Cannula placement was determined by visual identification under a light microscope of the guide cannula bore and/or lesion created by cannula insertion. Such points falling within the preoptic area boundary of Bregma - 0.3mm to Bregma +0.4mm and within 1mm above the MnPO were counted as hits. Data from misses regardless of experimental group were not included in any analysis.

### Steroid Treatments

Estradiol replacement paradigm followed our established protocol that mimics the natural physiological rise of E2 that occurs on the day of proestrus.^42, 63^ The replacement paradigm reliably reproduces (1) physiological levels of E2 equivalent to the day of proestrus^63^ and (2) changes in sleep-wake behaviors observed by intact cycling female rats.^40, 42^ Briefly, OVX females received 5 μg 17-β-estradiol benzoate in 5uL sesame oil (E2, SC; Sigma Aldrich, St. Louis, MO) followed by 10μg E2 in 10μL sesame oil 24 h later, or equivalent amounts (5µL/10µL) of sesame oil vehicle, through subcutaneous flank injections. The day following the last injection (Day 3 in the current set of experiments) is analogues to the day of proestrus. 17-β-estradiol benzoate is a synthetic ester of estradiol, which nonspecific steroidal esterases deesterify to produce biologically active E2 in vivo.^64, 65^ Of note, when testing the efficacy of the A_2A_ receptor agonist, CGS-2168, to induce sleep in the absence or presence of E2, a sub-physiological dose of E2 was administered (1.25µg and 2.5µg) in order to avoid the E2 mediated effects on sleep-wake behavior (see experiment 3 below).

### Data Acquisition, Sleep Scoring and Spectral Analysis

EEG and EMG data were collected at a sampling rate of 500Hz using the Ponemah Software (DSI Inc, St. Paul, Minn.). Digitized signal data were viewed offline using NeuroScore DSI v3.3.9 (Figure 1B; DSI Inc, St. Paul, Minn.). The EEG/EMG signals were parsed into 10 second epochs. A Fast Fourier transform (Hamming window, 4096 samples) was used to break down the EEG frequency bands (Delta (0.5-4Hz), Theta (4-8Hz), Alpha (8-12Hz), Sigma (12-16Hz), Beta (16-24Hz) Gamma (24-50Hz) and Total (0.5-50Hz)). The mean of the absolute value was calculated for the EMG data (bandpass 20-200Hz).

These data were exported to Matlab (Matlab R2015, Mathworks, Natick, Mass.) where vigilant states were automatically scored using a custom program developed in house (DJP) which has been shown to be ∼88-92% in agreement with hand scored traces from 3 independent scorers. In the automated program, scoring decisions were based on threshold levels of EEG delta power, theta power, the ratio of delta to theta, and the EMG activity. Data were normalized to the mean of the entire recording, then the median for each signal was used as a threshold (Supplementary Table 1). Artifacts were removed when data were 3 standard deviations greater than the mean of the surrounding 24 data points.

Finally, the scoring data was reimported into Neuroscore using a custom Matlab program courtesy of Dr. Michael Rempe (Whitworth University). Once in Neuroscore, visual inspection of the scored traces by an experimenter blinded to the treatment groups confirmed the automated scores and changes were made where necessary.

To further test whether E2 influences sleep homeostasis, NREM Slow Wave Activity (NREM-SWA; a marker of sleep homeostasis), was assessed via EEG spectral distributions of NREM sleep bouts. From Neuroscore, the EEG power spectra were calculated for NREM-SWA (0.5-4Hz), also referred to as delta power. Additionally, the frequency distribution of the EEG power spectral between 0.5-20Hz was calculated into 0.25Hz stepwise bins. Each power bin was normalized to the mean total power from the 24-hour baseline recording. Slow wave activity or the percentage of delta power over the total power was averaged into 1h bins during the light phase. For both slow wave activity and spectral power, artifact was removed if data were 3 standard deviations from the mean of the surrounding 24 data points.

### Experiments

#### Testing whether E2 influences sleep and sleep homeostasis in the light phase

To test whether E2 influences homeostatic sleep in the light phase, animals were randomly assigned to treatment arms in a double crossover design. The use of a within-animal blocking design mitigates the variance in the data resulting from differences in individual animal’s sleep-wake behavior. Adult female Sprague-Dawley rats (n=11) were OVX, implanted with EEG/EMG transmitters and randomly assigned to one of two sleep conditions: *ad libitum* sleep (AL) or sleep deprivation (SD). In the SD arm, animals were sleep deprived for the first 6 hours of the light phase on Day 3 and then allowed recovery sleep in the second 6 hours. The EEG/EMG for all groups started at Zt 0 and ended at Zt 12 (Light phase) on Day 3.

Once assigned to a sleep condition arm, animals were administered either oil vehicle or E2 according to our standard paradigm.^40, 42^ After Day 3 light phase recordings were collected, animals received a 7-day washout/rest period before crossing over into the other steroid/oil treatment. Again, recordings were collected on Day 3 of treatment. Following a second 7-day washout/rest period, all animals were crossed over to the opposite sleep condition arm. The steroid treatments and EEG/EMG recordings proceeded as in the first round.

Sleep deprivation was induced in the animal’s home cage between Zt 0-6. The animals were never removed or handled but were presented with novel objects that included wooden blocks, paper towel, and cotton balls to induce exploration. When the novel objects were no longer sufficient to maintain arousal, animals were gently stroked on the nape of the neck or back with a long cotton tip applicator. Finally, in the late stages of SD when the first two methods failed to prevent the initiation of sleep, it was necessary to gently rotate the home cage. Overall, the amount of accumulated Wake, NREM sleep and REM sleep was similar for both Oil and E2 treatment days where both groups were awake ∼88% of the SD period (Supplemental Figure S1). The majority of accumulated NREM sleep occurred in bouts that were less than 1 minute and most often less than 30 seconds (Supplemental Figure S1; inset). There was less than 0.1% of accumulated REM in either group (Data not shown).

Sleep-wake behavior was scored from the recorded EEG/EMG traces from Day 3 (Zt 0-12) of each treatment arm of the experiment. The degree of sleep pressure was assessed by established measures that included vigilant-state durations, bout length distributions, NREM-SWA across the phases, and NREM-SWA delta band power (0.5-4.5Hz). *A priori* comparisons between the Oil and E2 treatments under the specific sleep condition were made to test whether E2 affected sleep homeostasis. Additionally, the steroid treatment groups were collapsed and comparisons between AL and SD were made.

#### Testing whether E2 influences MnPO extracellular adenosine levels

Microdialysis collection during the light phase and HPLC analysis of interstitial fluid from the MnPO was used to investigate whether E2 altered levels of extracellular adenosine in both times of normal (AL sleep) and induced sleep need (SD). In one cohort, OVX animals were treated with either E2 (n=9) or Oil (n=10) and were allowed ad libitum sleep during the sample collection (AL cohort). Within this group, a small cohort of animals were fitted with telemeters to confirm that the AL cohort was accumulating sleep. In a separate cohort, OVX animals were treated with E2 (n=7) or Oil (n=7) and were subjected to 6 hours of sleep deprivation (SD) during the collection period. To maximize the likelihood of measuring differences between E2 and oil treated animals, the baseline measurements, which were used to normalize changes during the light phase collection period, were collected during the last hour of the wake phase when adenosine was expected to be at its peak.

Briefly, adult female rats were OVX and implanted with a microdialysis guide cannula targeted to the 1mm above the MnPO (see above). Following a 7-day recovery, the steroid treatment started with injections of either E2 or Oil vehicle. On Day 2, approximately 6 hours after treatment injections, the microdialysis probe (6kD membrane cutoff; Scipro inc., Sanborn, N.Y., model #MAB-9.14.1) was inserted into the guide cannula. The 7mm probe extended 1mm beyond the end of the guide cannula to rest within the MnPO. The surrounding tissue was allowed approximately 12 hours of probe acclimation without dialysis (∼Zt 22.5).

Prior to the start of dialysis, the probe was attached via polyethylene tubing to a 1ml Hamilton syringe (700 series, Hamilton, Reno, Nev.). The flowrate of the syringe was controlled by a BASi Bee pump attached to a Bee Hive controller (Bioanalytical Systems, Inc., West Lafayette, Ind.). The inserted dialysis probe was perfused at a rate of 1.167µL/minute with Ringer’s Solution (147mM NaCl, 4mM KCl, 1.4mM CaCl2, in distilled water). Probes were primed for at least 30 minutes before baseline collections began at Zt 23. During collection, animals could move freely about the cage and were provided food and water *ad libitum*.

Dialysis samples were collected in fractions of 20 minutes (23.3µL dialysate) for 7 hours. Upon collection, the dialysates were immediately frozen and stored at -20°C until LC-MS analysis. The first three fractions were collected within the last hour of the dark phase (Zt 23-0) and constituted the baseline of extracellular adenosine levels that was used to calculate the percent change in the light phase from Zt 0-6 (see below).

Probe placement was verified for each animal via light microscopy of Nissl stained sections. Placement was considered to be a hit if the probe/cannula fell within the MnPO as designated by the boundaries 0.48mm to -0.24mm from Bregma and 0.5mm right or left of the midline.^66^ For those animals designated as having correct placements extracellular adenosine levels were quantified.

To calculate the approximate *in vitro* probe recovery rate, free microdialysis probes (n=3) were inserted into a solution of 100nM adenosine (Tocris Biosciences, Bristol, U.K.) in Ringer’s Solution, and perfused with Ringer’s Solution at 1.167uL/min for 2 hours, with 20-minute dialysate fractions collected.

Adenosine quantification was performed by the Proteomics Core Laboratory in the Center for Vascular and Inflammatory Diseases at the University of Maryland, School of Medicine. Adenosine content in the collected fractions was quantified by liquid chromatography tandem-mass spectrometry by monitoring the transition pair of m/z 268.1/136.1 and quantified by plotting the area under the curve versus the known concentrations of the standards from a calibration curve. Analysis was performed on a Perkin Elmer (Waltham, Mass.) Qsight LX50 HPLC system and a QSight 210 triple-quadrupole mass spectrometer. The chromatographic solvents used were 0.1% formic acid in water (Solvent A) and 0.1% formic acid in methanol (Solvent B). The column was a YMC Triart 3µ C18, 2.1mm x 150mm operated at a flow rate of 400µl/min at 45°C. An isocratic separation was used with a solvent composition of 94% Solvent A and 6% Solvent B. The effluent from the column was introduced into the mass spectrometer by electrospray ionization in positive polarity and the transition pair of m/z 268.1/136.1 at unit mass resolution was used for detection of adenosine. The run time was 3 minutes. A stock solution of adenosine (Sigma) was prepared from dry powder and finally diluted in Solvent A to obtain a 6-point calibration curve that ranged from 5pg – 1500pg injected on column. The area under the curve (AUC) for the adenosine standards was plotted against their known concentrations to quantify the amount of adenosine in the experimental samples.

#### Testing whether E2 effects adenosine A_2A_ receptor activity

To test whether E2 attenuates A_2A_ receptor signaling in the MnPO, a highly selective A_2A_ agonist, CGS-21680 (CGS; Tocris Biosciences, Bristol, U.K.), was locally infused into the MnPO in the presence and absences of E2. Adult female Sprague-Dawley rats (n=10) were OVX and implanted with EEG/EMG transmitters and guide cannula. Following recovery, animals were randomly assigned to one of four treatment arms that consisted of subcutaneous steroid/Oil injections and local infusions of CGS or Vehicle. To reduce inter-animal variability, all animals served as their own controls and received all treatment (Oil/DMSO vehicle, E2/DMSO, Oil/CGS, and E2/CGS) in a random order. At the end of one treatment, animals were allowed a 7-day washout period before being randomly assigned to the next treatment.

To circumvent E2 suppression of sleep behavior, a sub-physiological dose of E2 that does not induce sleep suppression was administered (1.25µg and 2.5µg) according to the standard timing. Local MnPO microinfusions occurred at just prior to Zt 16 following the last E2 or Oil injection. The dummy stylets were removed and replaced by 33-gauge microneedles (PlasticsOne) that project 1mm past the end of the guide cannula. Infusion stylets were attached via polyethylene tubing to a 25μl Hamilton syringe (700 series, Hamilton). The flowrate of the syringe was controlled by a BASi Bee pump attached to a Bee Hive controller (Bioanalytical Systems). Infusions of 24nmol of CGS-21680 or equivalent volume of vehicle (3.89% DMSO in sterile saline) occurred over 10 minutes. At the end of the infusion, the needle was left in place for another 3-5 minutes to prevent back flow of the injection. The dose of CGS was previously reported to markedly increase sleep in male rats when infused into the lateral ventricle.^61^ Once the needle was removed and dummy stylets replaced, animals were returned to their home cages where EEG/EMG recordings were collected for the eight hours following the MnPO microinjection.

### Statistics

Results are expressed as means ± SEM. All statistical tests were conducted using the Graph Pad Prism program (San Diego, CA) on a PC. Statistical test and results are described in detail in the figure legends.

## Results

### E2 increased wake and decreased NREM sleep durations in recovery sleep but not during ad libitum sleep

Previous studies in female rats have demonstrated that E2 does not significantly influence unrestricted NREM sleep durations during the light phase unlike its marked effects in the dark^40, 42, 63, 67^ The current set of experiments sought to test whether E2 influenced homeostatic sleep pressure during (1) unrestricted light phase sleep (i.e. AL sleep) and (2) recovery sleep following a homeostatic challenge of 6 hours SD (Figure 2A). Compared to the oil control day, E2 significantly decreased the time spent in NREM sleep by ∼10% while increasing the time spent in wake during SD recovery sleep (zt 7-12) by ∼20%. As previously reported, there was no effect on NREM sleep or wake durations in AL sleep for the same 6-hour period (Figure 2B and C). Curiously, in AL sleep, E2 treatment significantly increased REM sleep by ∼17% from the oil control day. When the entire 12-hour light phase was analyzed, no significant differences in sleep-wake duration between E2 and control treatments were found (Supplementary Figure 2A).

**Figure 2.**
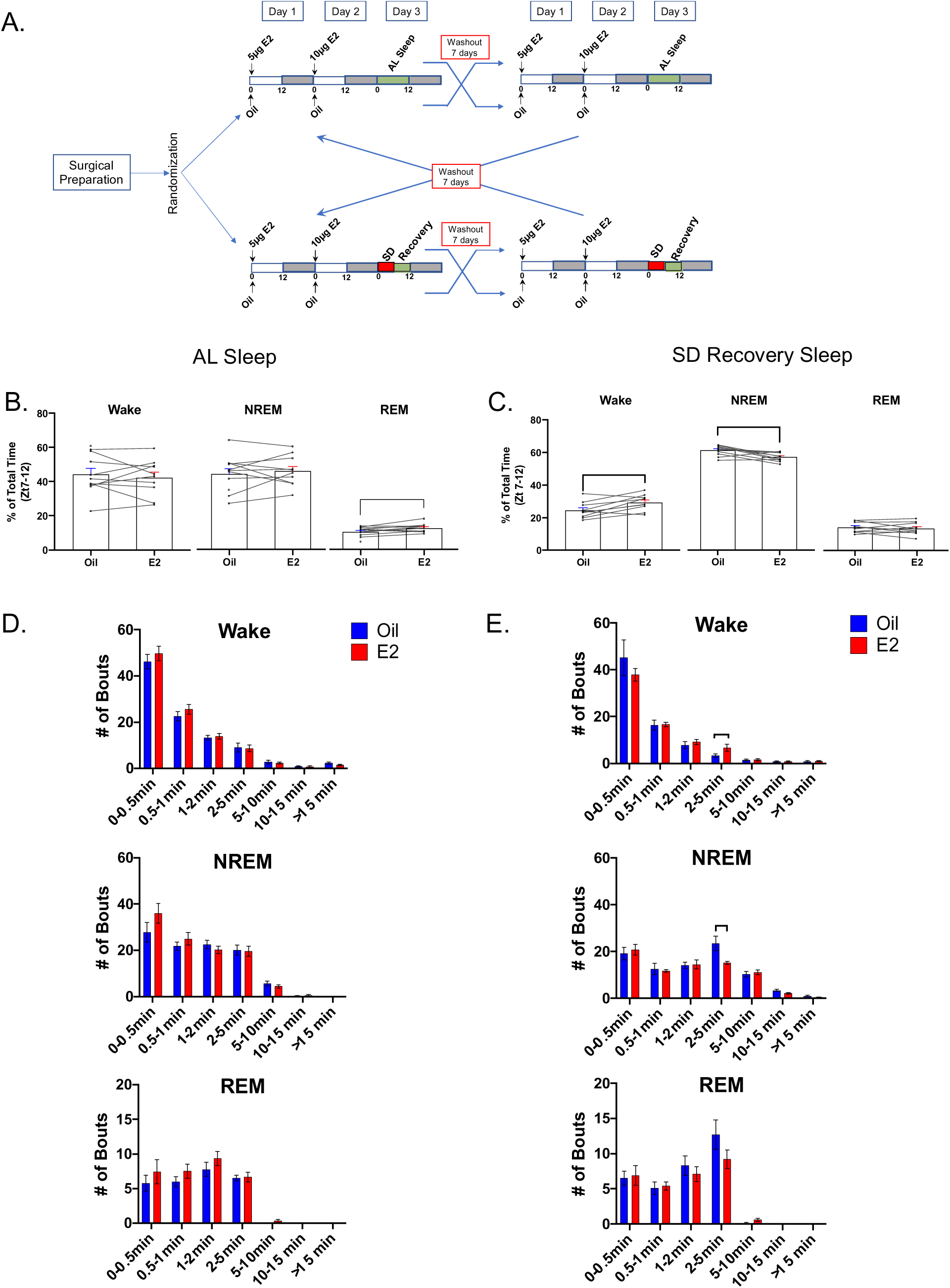
E2 increased wake and decreased NREM sleep durations in recovery sleep but not during AL sleep. **(A)** *Double cross-over experimental design.* Ovariectomized Sprague-Dawley rats (n=11) were implanted with transmitters and randomized to one of two sleep conditions: *ad libitum* sleep (AL) or sleep deprivation (SD). Once assigned to a sleep condition, animals were administered either oil vehicle or E2 according to our standard paradigm. At the end of the first recording period and 7 day washout period, the animals were crossed over to the other treatment but remained in the same sleep condition. At the end of the first cross over (and washout period), the animals were crossed to the other sleep conditions and treated as described above. Data are the mean±SEM. Individual lines represent paired measurements from the same animal. **(B)** *During AL sleep, E2 increased REM sleep but did not significantly affect Wake and NREM sleep.* When the second half of the Day 3 light phase (Zt 7-12) was analyzed for the time spent in each vigilant state, E2 did not significantly affect the percent of total time spent in Wake (Paired two-tailed t-test; t_(10)_=0.1665, p=0.8714) or NREM sleep (Paired two-tailed t-test; t_(10)_=0.3175, p= 0.7581). E2-treatment significant increased the percentage of REM sleep, albeit only by 1.2 fold, compared to oil controls (Paired two-tailed t-test; t_(10)_=2.566, *p= 0.0304). However, this difference in REM sleep between E2 and oil treatment was lost over the 12-hour light-phase (Supplemental data; Figure S2). Data are the mean±SEM. Individual lines represent paired measurements. **(C)** *During SD Recovery sleep, E2 increased Wake at the expense of NREM sleep.* When the period of SD Recovery sleep (Zt 7-12) was analyzed for the time spent in each vigilant state, E2 treatment significantly increased the percent of total time spent in Wake (Paired two-tailed t-test; t_(10)_=3.466, **p=0.0071), while significantly decreasing the percent of NREM sleep (Paired two-tailed t-test; t_(10)_=3.269, **p= 0.0097). The amount of REM sleep was unchanged (Paired two-tailed t-test; t_(10)_=0.7109, p= 0.4951). Data are the mean±SEM. **(D, E)** *In SD Recovery Sleep, E2-treatment significantly influenced bout length distribution.* Wake, NREM sleep and REM sleep bouts were binned into 7 time intervals representing short (bouts lasting less than 1minute), medium (bouts between 1-5 minutes) and long (bout greater than 5 mins) durations. Similar to the time spent in each vigilance state, a Wilcoxon matched pairs signed rank test demonstrated that E2 did not affect the bout distribution of any state in AL sleep on Day 3. However, in SD Recovery sleep, the Wilcoxon matched-pairs signed rank test revealed that E2 increased the number of 2-5 minute Wake bouts (Holm-Šídák correction for multiple comparisons, *p= 0.0409), while decreasing the number of NREM sleep bouts in the same interval (Holm-Šídák correction for multiple comparisons, *p= 0.0403). Data are the mean±SEM.

Analysis of bout length distributions for the AL and SD recovery sleep period revealed that E2 specifically influenced SD recovery wake and NREM bouts that were between 2-5 minutes with no effect in AL sleep (Figure 2 D, E). Overall, vigilant state bout totals were unchanged by E2, except for an ∼20% increase over oil controls in AL REM sleep (Table 1).

**Table 1:**
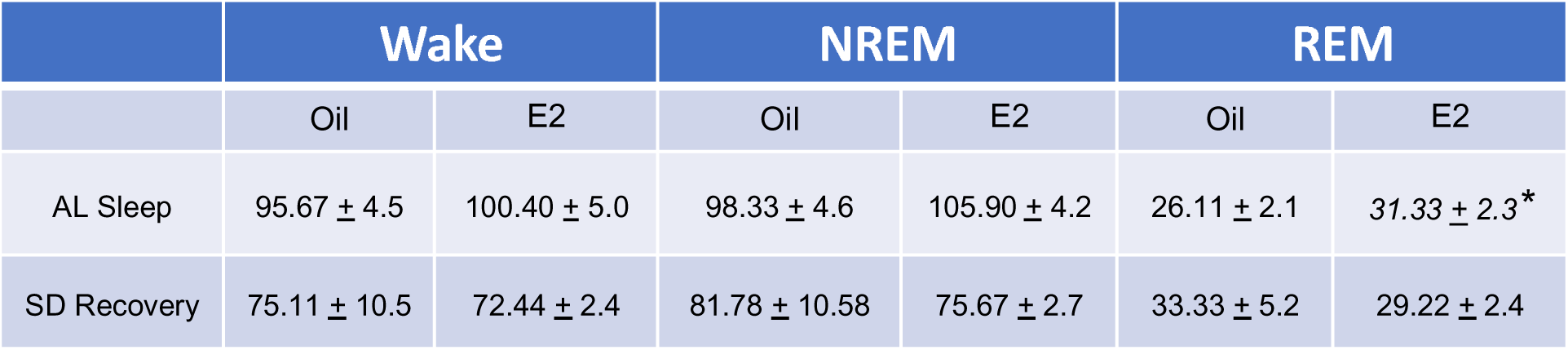
Bout totals in AL and SD Recovery Sleep during (Zt 7-12). In either AL sleep or SD Recovery Sleep, E2 treatment did not significantly affect the total number of Wake or NREM bouts when compared to the oil treatment within the same sleep group. In AL sleep, E2 treatment increased the number of REM bouts compared to oil (Paired two-tailed t-test; t_(10)_=3.271, *p=0.0113), but not in SD Recovery. Data are the mean±SEM.

### E2 decreased SWA in AL sleep but not in recovery sleep

To further explore E2 effects on sleep homeostasis, the amount of SWA (0.5-4.5Hz) and spectral power distributions during light phase NREM sleep bouts were compared to Oil controls (Figure 3 A-D). Analysis of the NREM-SWA per hour revealed that E2-treatment significantly decreased the hourly amount of SWA during NREM bouts compared to Oil in AL sleep but not SD recovery sleep (Figure 3A, B). In AL sleep the decreased NREM-SWA was also seen across the 12-hour light phase (Supplemental Figure S2).

**Figure 3.**
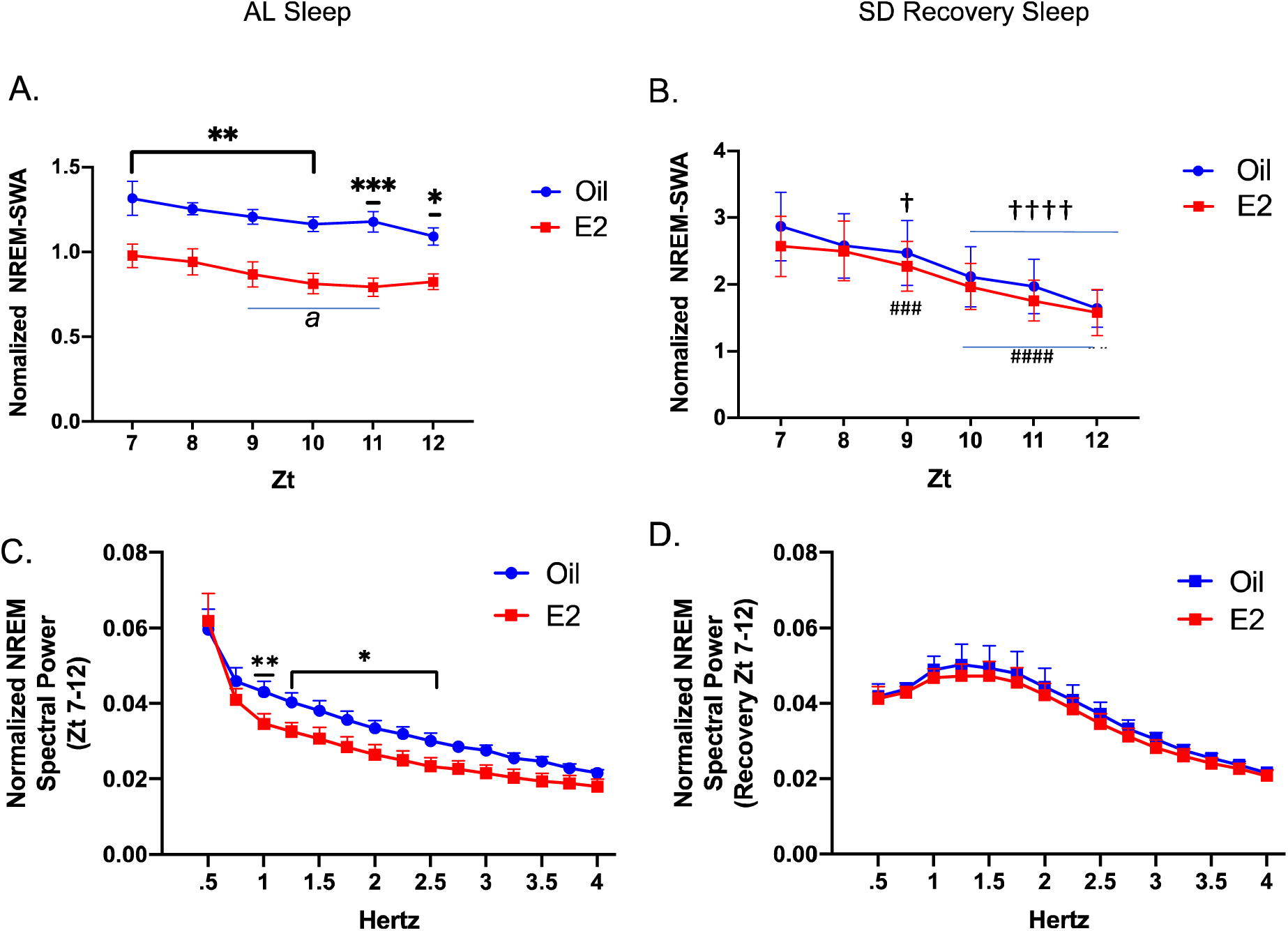
E2 decreased NREM-SWA in AL sleep but not SD Recovery Sleep. (A,B) *Hourly totals of SWA in NREM sleep bouts.* For all NREM bouts on Day 3 (Zt 7-12), the EEG power spectra in the 0.5-4Hz range (also known as delta power) was calculated and normalized to the mean total power from the 24-hour baseline recording. The normalized NREM-SWA was calculated for each hour from Zt 7-12. In AL sleep, E2 significantly decreased the amount of NREM-SWA compared to oil treatment across the 6 hour period (Repeated measure two-way ANOVA; main effect of E2, F_(1,48)_=77.15; p<0.0001). A post hoc multiple comparison revealed significant differences at each of the hours (Šídák’s multiple comparison test; Zt 7-10, **p<0.01; Zt 11, ***p<0.001; Zt 12 *p<.05; see supplemental Table 2 for adjusted P values and confidence intervals). Similarly, across the entire 12-hour light phase E2 significantly reduced NREM-SWA (Supplementary Fig S2). Interestingly, there was a main effect of time (F_(5,48)_=3.143, p<0.0156) where a multiple comparison post-hoc test revealed significant differences only in the E2-treatment group (Sidak’s multiple comparison test; Zt 7 vs. Zt 9, Zt 10 and Zt 11, p<0.05 *a denotes difference from Zt 7*; see supplemental Table 2 for adjusted P values and confidence intervals) suggesting that E2 treatment may contribute to a more rapid resolution of sleep need. There was no interaction between E2 and Zt (F_(5,48)_ = 0.1854; p=0.9668). In recovery sleep, NREM-SWA was not significantly different between E2 and oil treatments (Repeated measure two-way ANOVA; effect of E2, F_(1,48)_=0.0843; p=0.7758); however there was a main effect of time (F_(5,48)_=34.33, p<0.0001), where a multiple comparison post-hoc test revealed significant differences in both the E2 and oil treatment groups (Sidak’s multiple comparison test: Oil: Zt 7 vs. Zt 9, ^†^p=.0391 and Zt 7 vs. Zt 10, Zt 11, Zt 12, ^††††^p<0.0001 and E2: Zt 7 vs. Zt 10, ^###^p=0.0004; Zt 7 vs Zt 11, Zt 12 ^####^p<0.0001 see supplemental Table 2 for adjusted P values and confidence intervals). Again there was no interaction between E2 and Zt (F_(5, 48)_ = 0.3795; p=0.8612). Data are the mean±SEM. (C, D) *Spectral Power Distribution in NREM sleep.* In AL sleep, analysis of the spectral power distribution for NREM sleep revealed that E2-induced reductions in spectral power were limited to the lower frequency delta range in both the first (Zt 0-6) and second (Zt 7-12) half of the light phase, while there was no change in the spectral distribution of NREM sleep during SD recovery (Supplemental Figure S3 A-C). Comparison of spectral distribution within the Delta frequency range (0.5-4 Hz) from Zt 7-12 between E2 and oil treatments revealed a main effect of E2 (F_1,120_= 94.85, p<0.0001) and frequency (F_14,120_=18.49, p<0.0001) and no interaction between the two factors (F_14,120_=1.310; p=0.2112) in AL sleep, while in SD recovery sleep there was no significant differences between E2 and oil treatments. A multiple comparison post-hoc test of the delta range spectral distribution revealed significant differences between E2 and oil treatment in the 1Hz-2.5Hz range (Sidak’s multiple comparison test; **p<0.01 and *p<0.05; see supplemental Table 2 for adjusted P values and confidence intervals). Data are the mean±SEM.

To address whether sleep need was decreasing across the AL sleep phase, the amount of NREM-SWA was analyzed as a function of time for the given period (Zt 7- 12). E2 but not oil treatment demonstrated a significant decrease at Zt 9, 10 and 11 compared to Zt 7 (Figure 3A; *a* denotes Zt points that are significant from Zt 7). Moreover, the distribution of spectral power revealed that E2 treatment significantly decreased delta power in the lower frequency range (1-2.5 Hz) compared to Oil during AL sleep in both the first (Zt 0-6) and second half (7-12) of the light phase but not in SD recovery sleep (Figure 3C, D and Supplemental Figure S3).

### E2 induced a greater magnitude of change in SWA during recovery sleep

To test whether E2 affected the recovery from SD, NREM-SWA and NREM spectral power distributions from E2 and oil control days were compared between the AL and SD recovery periods (Zt 7-12; Figure 4 and 5). As expected, NREM-SWA activity significantly increased during the SD recovery phase compared to the same time-period during AL sleep for both Oil and E2 treatment (Figure 4A, B). Additionally, NREM-SWA dissipated over the course of the recovery period where SWA during the last 3 hours of SD-recovery (Zt 10-12) was significantly less than the first hour of SD-recovery for both Oil and E2 treatment (4A, B). Analysis of the percent change in SWA between the SD recovery and AL sleep periods revealed that E2 treatment resulted in a significantly greater magnitude of change with SD compared to oil (Figure 4 C). Similarly, the distribution of spectral power revealed greater power in the lower frequency ranges (Oil: 1.5-2.25hz and E2: 1-2.5hz) during SD recovery compared to AL sleep (Figure 5 A, B). Analysis of the magnitude of change (% change) in SWA between the SD recovery and AL sleep periods revealed a main effect of E2 treatment compared to Oil (Figure 5C). Furthermore, an Area Under the Curve (AUC) analysis of the magnitude of change suggested that E2 induced an ∼2-fold increase over oil (Figure 5D).

**Figure 4.**
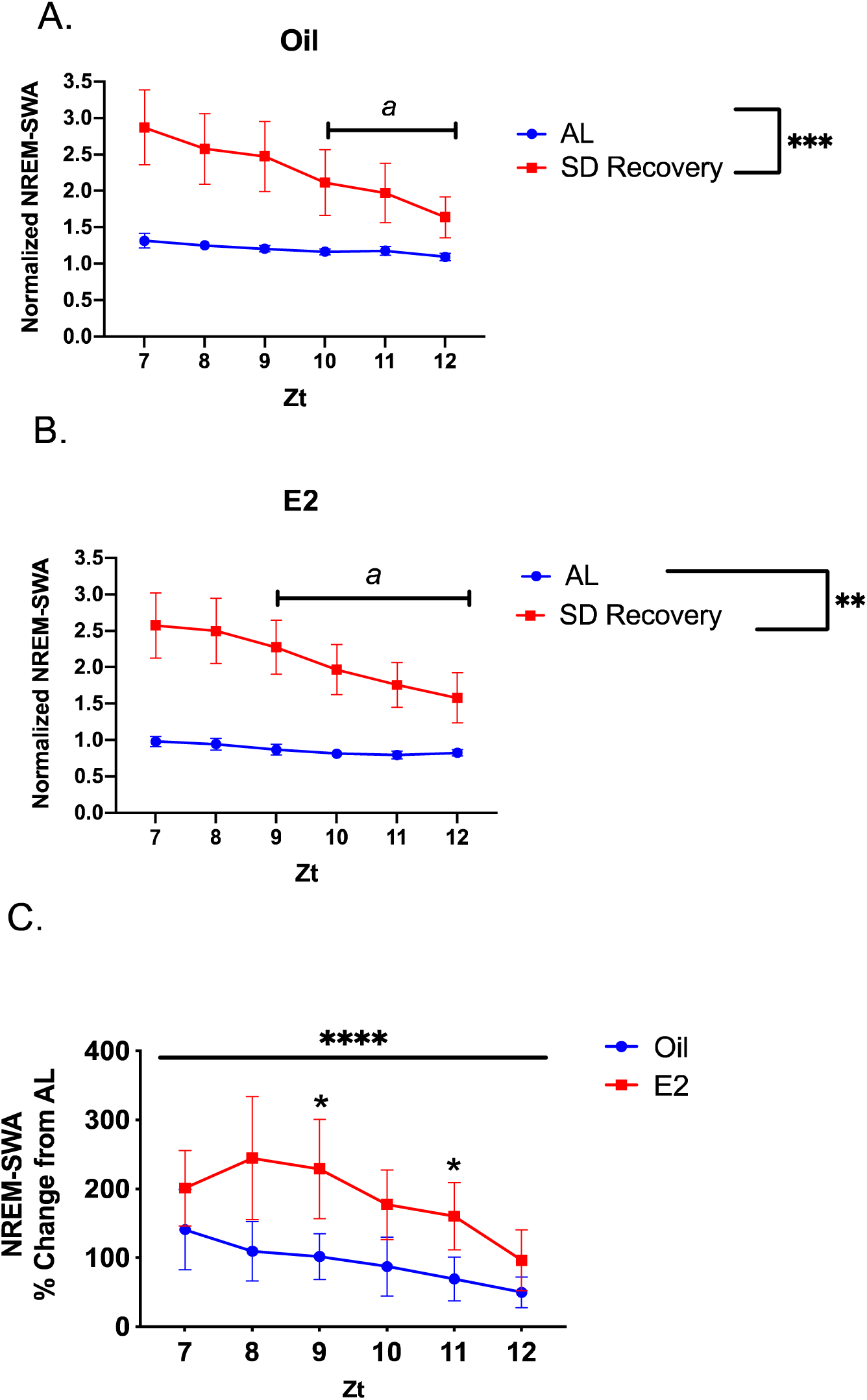
E2 induced a greater magnitude of change in SWA during recovery sleep. (A) *Comparison of NREM-SWA in AL and SD recovery sleep following oil treatment.* As expected, compared to the undisturbed sleep condition in the AL group, NREM-SWA activity increased significantly in the recovery period that immediately followed 6-hours of total sleep deprivation. The comparison of NREM-SWA between AL and SD recovery across the 6-hour period revealed main effects for both factors (Repeated measure two-way ANOVA; main effect of sleep condition, F_1,8_ = 40.94, p=0.0002; main effect of Zt, F_5,40_= 3.969, p=0.0051; ****represents the main effect of sleep condition), as well as a significant interaction between the two factors (F_5,34_= 5.733, p=0.0006). A multiple comparison post-hoc comparison of Zt 7 to the other hours of the respective sleep condition revealed that only SD recovery sleep exhibited a significant decrease in NREM-SWA across time (Sidak’s multiple comparison test; Zt 7 vs Zt 10, Adjusted **p=0.0048; Zt 7 vs Zt 11, Adjusted ***p=0.0006; Zt 7 vs Zt 12, Adjusted ****p<0.0001; a denotes significant differences from Zt 7). (B) *Comparison of NREM-SWA in AL and SD recovery sleep following E2 treatment.* Similar to the control condition, NREM-SWA activity was significantly increased in the SD recovery period compared to the AL sleep period under E2-treatment. The comparison of NREM-SWA between AL and SD recovery across the 6-hour period revealed main effects for both factors (Repeated measures two-way ANOVA; main effect of sleep condition, F_1,8_=12.17, p=0.0082; main effect of Zt, F_5,40_= 26.35, p<0.0001; **represents the main effect of sleep condition), as well as a significant interaction between the two factors (F_5,34_=13.48, p<0.0001). The multiple comparison post-hoc between Zt 7 and the other hours of the respective sleep condition revealed that with E2 treatment again only SD recovery sleep exhibited a significant decrease in NREM-SWA across time (Sidak’s multiple comparison test: SD: Zt 7 vs 9 Adjusted *p=0.0106; Zt 7 vs 10, 11 and 12, Adjusted ****p<0.0001; *a denotes significant differences from Zt 7 for SD Recovery*). (C) *Comparison of the percent change of NREM-SWA between oil and E2 treatment.* To determine whether E2 affected the magnitude of change from a baseline condition following a significant homeostatic challenge, the percent change of hourly NREM-SWA between SD recovery and AL sleep was calculated for both oil and E2. Comparison of the resulting differences revealed that E2 had a significantly greater magnitude of change compared to oil (Repeated measures two-way ANOVA; main effect of E2, F_1,42_=25.00, ****p<0.0001). A post-hoc comparison additionally demonstrated significant differences at Zt 9 and 11 (Šídák’s multiple comparison test, Adjusted p values: *p= 0.0264 and 0.0414, respectively. There was no effect of Zt or significant interaction. Data are the mean±SEM.

**Figure 5.**
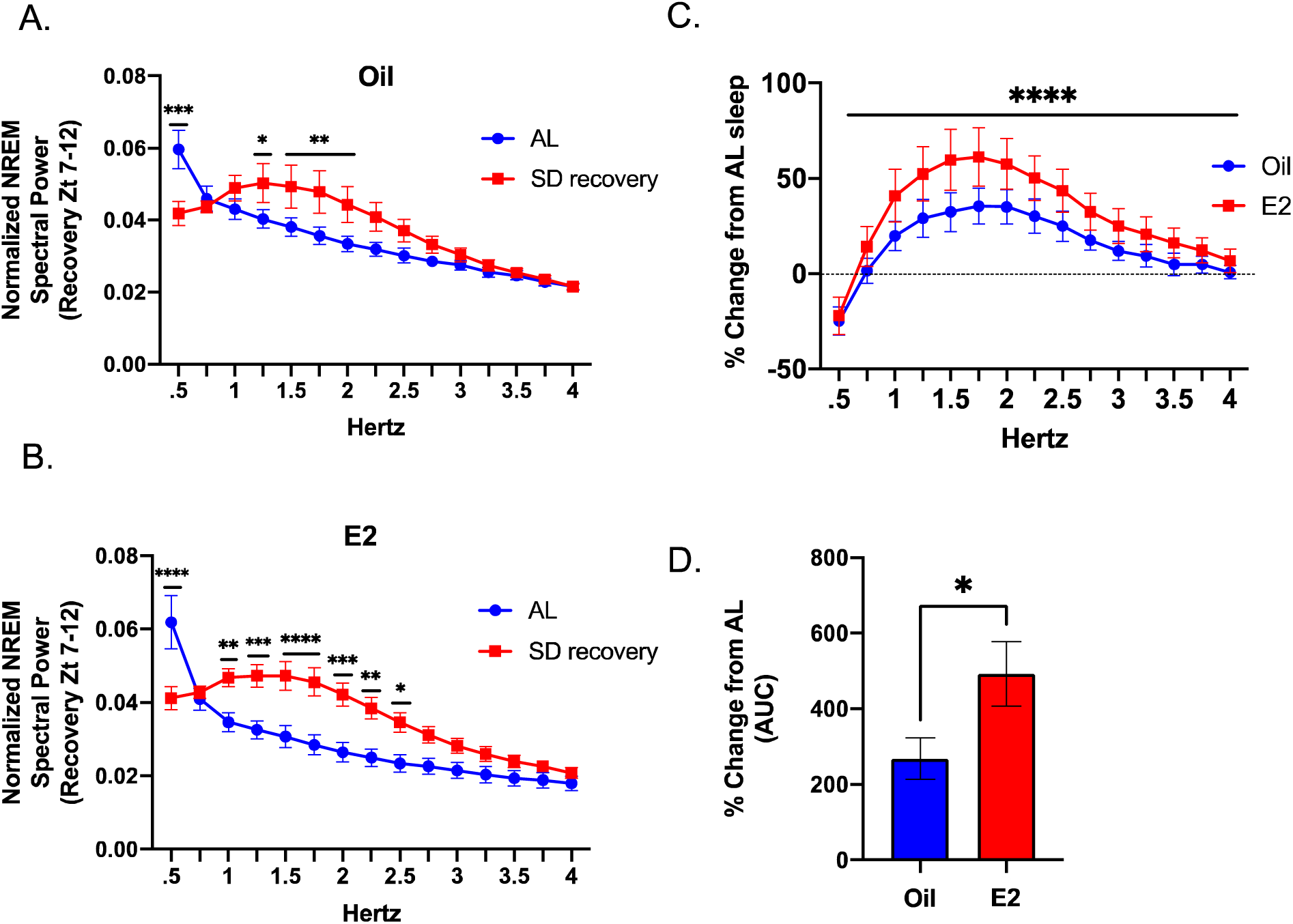
E2 induced a greater magnitude of change in the spectral distribution of delta power during recovery sleep. **(A)** *Comparison of the spectral distribution in delta frequencies from AL and SD recovery sleep following oil treatment.* Given that the 6-hour sleep deprivation significantly increased NREM-SWA (Figure 4), it was highly anticipated that a spectral distribution of the low frequency delta bands would exhibit a marked increase in power compared to the spectral distribution in AL sleep. The comparison of spectral distributions following oil treatment between AL and SD recovery revealed main effects for sleep condition and frequency (Repeated measures two-way ANOVA; main effect of sleep condition, F_1,120_= 23.59, p<0.0001; main effect of Zt, F_14,120_=12.69, p<0.0001) as well as a significant interaction between the two factors (F_14,120_=6.190, p<0.0001). A multiple comparison post-hoc test revealed that, overall, SD recovery sleep had significant increases in the 1.25Hz-2.0Hz range. (Šídák’s multiple comparison test, Adjusted p values: 0.5 Hz, ****p<0.0001; 1.25Hz, *p=0.0202; 2.5Hz, **p=0.0049; 2.75Hz, **p=0.0015; 2Hz, **p=0.0070). **(B)** *Comparison of the spectral distribution in delta frequencies from AL and SD recovery sleep following E2 treatment.* Like oil treatment, the spectral distribution of the low frequency delta bands exhibited a marked increase in power across the lower ranged frequencies compared to the spectral distribution in AL sleep. The comparison of spectral distributions following oil treatment between AL and SD recovery revealed main effects for sleep condition and frequency (Repeated measures two-way ANOVA; main effect of sleep condition, F_1,120_= 75.44, p<0.0001; main effect of Zt, F_14,120_=17.53, p<0.0001) as well as a significant interaction between the two factors (F_14,120_=7.701, p<0.0001). A multiple comparison post-hoc test revealed that, overall, SD recovery sleep had significant increases in the 1.0Hz-2.5Hz range. (Šídák’s multiple comparison test, Adjusted p values: 0.5 Hz, ****p<0.0001; 1.0Hz, **p=0.0076; 1.25Hz, ***p=0.0005; 1.50Hz, ****p<0.0001; 1.75Hz, ****p<0.0001; 2Hz, ***p=0.0001); 2.25Hz, **p=0.0018; 2.50Hz, *p=0.0196). **(C)** *Comparison of the percent change of the delta spectral distribution between oil and E2 treatment.* To determine whether E2 affected the magnitude of change from a baseline condition following a significant homeostatic challenge, the percent change of power in each delta band frequency between SD recovery and AL sleep was calculated for both oil and E2. Comparison of the resulting differences revealed that E2 had a significantly greater magnitude of change compared to oil (Repeated measures two-way ANOVA; main effect of E2, “F _1,120_ = 50.46, ****p<0.0001). (D) *Area under the curve comparison (AUC) of the overall magnitude of change.* An AUC analysis of the resulting precent change in (C) was used to assess the overall differences in change between oil and E2 and demonstrated that E2 treatment resulted in a significantly greater magnitude change in delta power compared to oil (Paired two-tailed t-test; t_(7)_=4.468, *p=0.0430).

### E2 increased extracellular adenosine content in the MnPO of adult female rats

Building on the finding that E2 decreased sleep behaviors and architecture that are characteristic of homeostatic sleep pressure, we next tested whether E2 decreased extracellular concentrations of adenosine in the POA via microdialysis. Animals were prepped and treated according to timeline in Figure 6A. The microdialysis probe placements within the MnPO are represented in Figure 6B. Based on the mean *in vitro* probe recovery of 12%, the average extracellular baseline adenosine concentration in the MnPO of females was calculated to be ∼ 203nM, which is near the range of reported values of 50-200nM.^68^

**Figure 6.**
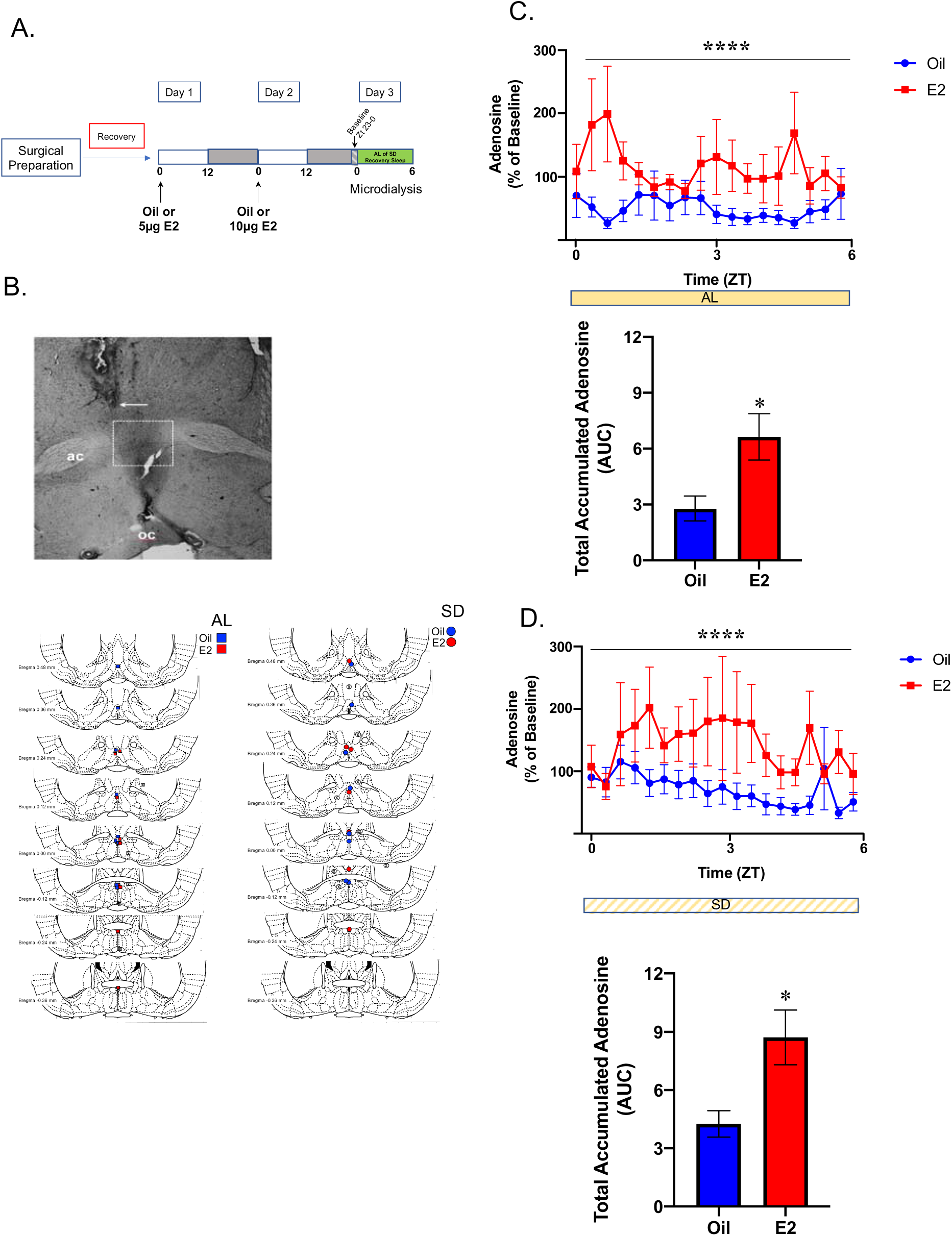
Microdialysis of MnPO extracellular fluid and adenosine content analysis. (A) *Experimental timeline*. Two separate experimental cohorts of Sprague-Dawley rats were OVX and implanted with microdialysis guide cannula targeted to the MnPO. The first experimental cohort was used to investigate the effects of E2 on extracellular adenosine of the MnPO. Following recovery from surgery, animals in the first cohort were treated with E2 (n=8) or oil vehicle (n=10) on two consecutive days prior of microdialysis. On Day 3 the collection of dialysates began in the last hour of the dark phase. These sampled served as the normalization baseline. For the remainder of the experiment, animals were allowed spontaneous sleep (AL sleep) during sample collection. The second experimental cohort (E2; n=7 and oil; n=7) was treated as described except during the sample collection the animals were subjected to the total sleep deprivation protocol. Of note, an a priori power analysis determined that to achieve greater than 80% power, samples sizes of n=10 were required. Throughout the course of both experiments, animals were either lost or cannulae were misplaced. In the case of the later the samples were not analyzed. A posthoc power analysis determined that the resulting sample sizes achieved 77% power in the AL sleep cohort and 75% in the SD cohort. (B) Cannula Placement. *Top.* Representative photomicrograph of the cannula and microdialysis probe placement. Probe placement was determined visually using neutral red staining. The arrow demarcates the end of the guide cannula from where the dialysis probe extended 1mm. The box represents the area of the MnPO. *Bottom.* Maps of the probe placements (including misses) for the AL and SD sleep cohorts. Samples were analyzed if the probe placement fell within the preoptic area boundary defined by Bregma -0.3mm to Bregma +0.4mm and within with boundary of the MnPO (as marked by the box in the photomicrograph above). *Blue symbols, oil; Red symbols, E2; X-mark, misses.* Adenosine levels were collected by microdialysis (6kd-cutoff probes) and analyzed by HPLC-Mass Spectrometry. Adenosine levels were normalized to each animal’s baseline adenosine collected from Zt 23-Zt 0. (C) *AL Sleep Cohort. Top.* Extracellular adenosine levels following E2 treatment were significantly elevated across the 6-hour collection compared to oil (Two-way ANOVA, main effect of E2; F_1,302_=35.44 ****p<0.0001). *Bottom.* Area under the curve analysis demonstrated that the total levels of extracellular adenosine were markedly increased following E2 treatment compared to oil (Two-tailed t-test; t_(17)_=2.832, p=0.0115). (D) *SD Cohort. Top.* Similar to undistributed sleep, extracellular adenosine levels following E2 treatment were significantly elevated across the 6-hour of sleep deprivation compared to oil (Two-way ANOVA, main effect of E2; F_1,221_=28.41, ****p<0.0001). *Bottom.* Area under the curve analysis demonstrated that the total levels of extracellular adenosine during sleep deprivation were markedly increased following E2 treatment (Two-tailed t-test; t_(12)_=2.842, p=0.0148). Comparing the AUC calculations across the four treatment groups suggested that SD did increase total adenosine content in both the oil and E2 treated groups compared to AL sleep; however, the increase did not reach statistical significance (Supplemental Figure S4). This could be due to the experimental variability between the AL and SD cohorts which were run as separate experiments. Data are the mean±SEM.

Extracellular adenosine content was significantly greater in E2 treated females compared to the Oil controls during the first 6 hours of light phase AL sleep (Figure 6C). An area under the curve (AUC) analysis over the 6 hours revealed that total accumulated adenosine significantly increased by ∼2-fold in E2 treated females compared to Oil controls (Figure 6C). Similarly, extracellular adenosine content was significantly greater in E2 treated females compared to the Oil controls during sleep deprivation that occurred over the first 6 hours of the light phase. An area under the curve (AUC) analysis for this SD period revealed that total accumulated adenosine was significantly increased by ∼3-fold in E2 treated females compared to Oil controls (Figure 6D).

Comparing the AUC calculations across the four treatment groups demonstrated that SD did increase total adenosine content in both the oil and E2 treated groups compared to AL sleep; however, the increase did not reach statistical significance (supplemental Figure S4). This could be due to the experimental variability between the AL and SD cohorts which were run as separate experiments. Finally, when *ad libitum* sleep-wake durations were analyzed in a subset of animals undergoing microdialysis and implanted with transmitters, E2 and Oil treated animals had equivalent amounts of light phase sleep (NREM and REM) and wake (like the results in Figure 2B), despite their differing adenosine levels similar (Data not shown).

### E2 treatment attenuated the sleep promoting actions of an adenosine A_2A_receptor agonist in the MnPO

Given that adenosine is widely accepted as both a molecular marker and mediator of sleep pressure,^56, 59, 69^ the finding that E2 increased MnPO adenosine content while reducing NREM-SWA presented an interesting paradox which suggested that E2 may be working to attenuate adenosine signaling and thus dampen the detection of homeostatic sleep pressure. To test this prediction, we investigated whether A_2A_R activation via local infusion of CGS, a highly specific agonist, induced sleep and decreased NREM-SWA in females with and without E2 (Figure 7). Animals were prepped and treated according to timeline in Figure 7A. The cannula placements for drug infusion into the MnPO are represented in Figure 7B.

**Figure 7.**
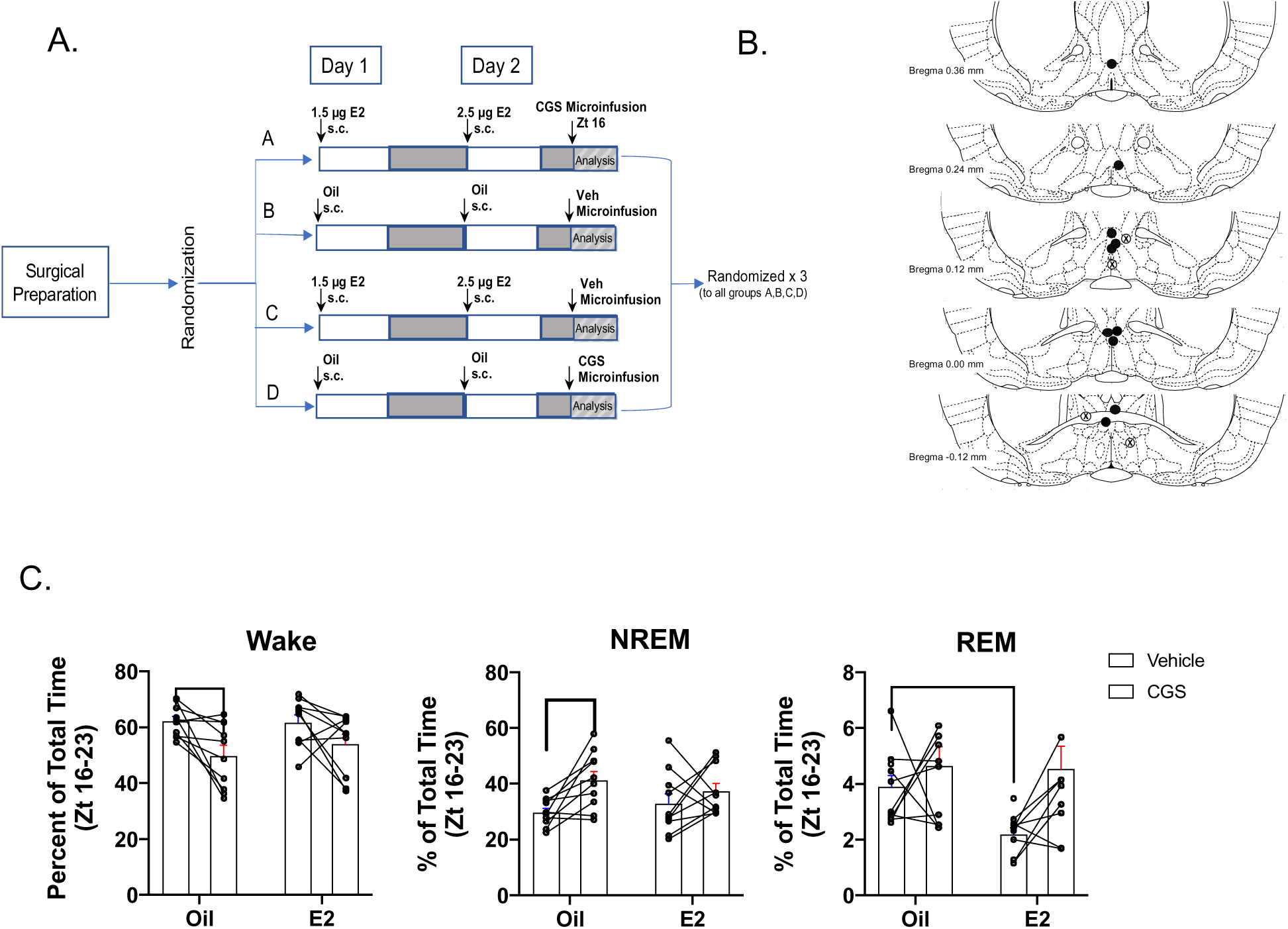
A2A Agonist CGS-21680 decreased wake and increases NREM sleep following oil but not E2 treatment. *Experimental timeline*. Sprague-Dawley rats were OVX, fitted with telemeters and implanted with infusion guide cannula targeted to the MnPO (n=10). Following recovery from surgery, animals were randomly placed into one of four treatment arms where they received 2 subcutaneous injections of either oil or a subthreshold dose of E2 (that does not affect sleep-wake behavior) and at Zt16 on Day 2 infusion into the MnPO of either the A2A agonist CGS-21680 (CGS; 24nmol) or DMSO vehicle. The EEG/EMG recordings began immediately after the MnPO infusions. At the end of the recording period the animals were allowed a 7-day washout period before being randomly assigned to another treatment arm. This protocol was repeated until all animals had been run through each treatment arm. The EEG/EMG traces from Zt 16-0 were scored and analyzed changes for changes in sleep-wake states. As CGS was predicted to increase NREM sleep at the expense of wake, the dark phase was chosen to maximize the probability of detecting changes in these two vigilant states. (B) *Cannula Placement.* Cannula placement was determined visually using neutral red staining. Representation of guide cannula placements (including misses). Placements were deemed a hit when they fell within the preoptic area boundary defined by Bregma -0.3mm to Bregma +0.4mm and within with boundary of the MnPO (as marked by the box Figure 5B). *Black circles= hits; Circled X=misses.* (C) Effects of CGS on Wake, NREM and REM sleep in the presence and absence of E2. Local infusion of CGS into the MnPO significantly decreased Wake following oil treatment, but not in the presence of E2 (Repeated measures two-way ANOVA; main effect of CGS F_1,9_=15.16; p=0.0037; no effect of E2; F_1,9_=0.2983, p=0.5982; Šídák’s multiple comparison test, Adjusted p value *p=0.0474). The decrease in Wake was accompanied by an increase in NREM sleep following oil treatment only (Repeated measures two-way ANOVA; main effect of CGS F_1,9_=8.822; p=0.0157; no effect of E2; F_1,9_=0.0178, p=0.8969; Šídák’s multiple comparison test, Adjusted p value *p=0.0312). Curiously, REM sleep showed a significant effect of E2 but not CGS (Repeated measures two-way ANOVA; main effect of E2 F_1,9_=5.576; p=0.0425; no effect of CGS; F_1,9_=2.233, p=0.1693; Šídák’s multiple comparison test, Adjusted p value *p=0.0.0246). Individual lines represent paired measurements from the same animal. Data are the mean±SEM.

In females treated with Oil, local infusion of the CGS into the MnPO significantly reduced wake and increased NREM sleep in the dark phase by ∼20% compared to vehicle infusion (Figure 7C,). This effect is similar to those reported in male rats.^61^ However, when CGS was infused in the presence of E2 treated animals, the CGS- induced increase in NREM sleep and decrease in wake was significantly attenuated (Figure 7C,). The agonist did not have a significant effect on REM sleep with or without E2 (Figure 7C). Of note, the combination of the lower E2/Vehicle did not significantly change wake or NREM but did decrease REM sleep compared to Oil/Vehicle controls.

## Discussion

The current set of findings provide evidence that E2 is involved in altering homeostatic sleep. Our previous work suggested that E2 might lower homeostatic need.^42, 63, 67^ In the current study, we investigated estrogenic effects on specific behavioral, electrophysiological and neuromodulatory markers of sleep homeostasis in the light-phase. In undisturbed light-phase sleep, E2 markedly reduced NREM-SWA, a marker of homeostatic need, in the absence of behavioral sleep changes. In contrast, in the recovery period following total sleep deprivation, E2 reduced the time spent in NREM, but the NREM-SWA was equivalent to the control treatment. However, E2 induced a greater magnitude of change in SD recovery sleep from baseline AL sleep. This finding suggests that E2 may expand the homeostatic set-point for sleep, allowing for a greater build-up in sleep pressure before initiating a response. Surprisingly, extracellular adenosine in the MnPO markedly increased in AL and SD recovery sleep conditions following E2-treatment. Although increases in extracellular adenosine are reliably linked to sleep induction, the finding that E2 attenuates A_2A_ receptor actions in the MnPO further suggests a possible mechanism through which E2 blocks the actions of adenosine and further contributing to decreasing sleep need. Taken together, the current sent of findings provide evidence that E2 may be working to increase the homeostatic set-point for sleep allowing for a decrease in sleep propensity despite periods of increased wake duration.

### Sleep Homeostat

A general consensus is that mammals require a set amount of sleep over a given time period and the brain is capable of sensing this sleep pressure or need. The brain’s intrinsic measurement of sleep pressure is referred to as the homeostatic sleep system.^53^ As the name implies, the sleep homeostat governs the amount of sleep needed after a given period of wake to maintain homeostasis. The existence of a sleep homeostat is best exemplified by the need for recovery sleep following periods of sleep deprivation.^51, 52, 70^ While the mechanism and/or circuitry of the sleep homeostat remains elusive, NREM-SWA and NREM sleep duration are characteristic hallmarks of sleep need.^56, 57^ Increases in the intensity or amount of NREM-SWA are proportional to the amount of prior wake time, while time spent in NREM sleep will dissipate the concentration of SWA. Given these dynamic changes that are proportional to sleep loss/gain, NREM-SWA is widely used as a reliable marker of sleep homeostasis.

### Estradiol and Sleep Homeostasis

In rodents, the absence of circulating gonadal steroids, eliminate sex differences in sleep behavior and architecture^40, 42, 71^, suggesting that sex differences in sleep are primarily dependent on circulating sex steroids. In adult female rats, endogenous and exogenous E2 markedly suppresses dark-phase NREM sleep; however, findings as to whether E2 significantly affects light-phase sleep are mixed.^62, 40, 72, 73, 42, 41^ One explanation for the mixed findings of E2 effects on light-phase sleep is the varied steroid-replacement paradigms and variability of endogenous steroidal milieus across animals. In our previous work, we found that Day 3 of the E2-replacement paradigm reliably mimics proestrus reproductive behaviors,^74, 75^ sleep,^40, 42^ and serum E2 levels.^63^ Thus, using this replacement paradigm which allows for a more standardized physiological level of circulating E2, here, we investigated whether E2 influenced markers of homeostatic sleep in the light phase.

Like the proestrus light-phase findings in cycling females,^41, 62^ E2-replacement significantly reduced the amount of NREM-SWA as well as decreased the power densities of low frequency delta bands across the 12-hour light phase. This decrease occurred in the absence of changes in NREM sleep times suggesting that E2 significantly reduced homeostatic sleep need compared to controls despite spending equivalent times in NREM sleep. To further assess E2 effects on homeostatic sleep regulation, the NREM-SWA response to 6-hours of total sleep deprivation was measured. Typically, sleep deprivation invariably leads to an increase in SWA during recovery sleep. If E2 reduces homeostatic sleep need, then a likely prediction is that SD-induced increases in NREM-SWA will be reduced following E2 treatment compared to control. However, the current findings did not entirely support this prediction. In the recovery phase, the intensity and power distribution of NREM-SWA between E2 and control treatment was not different even though E2 treatment significantly increased the time spent in wake at the expense of NREM sleep. Nevertheless, the E2-induced increase in wake following a significant homeostatic sleep challenge further suggests that sleep drive is diminished compared to oil controls. Moreover, the E2-induced increase in wake during SD recovery was not due to increased fragmentation of sleep, but rather an increase in wake bouts and a decrease in NREM bouts in the 2-5 minute range. These findings taken together further supports the assertion that E2 is damping the homeostatic sleep drive and allowing for stable consolidated wake bouts.

Comparing the magnitude of changes in NREM-SWA between AL and SD recovery sleep further revealed characteristics of E2 effects on sleep homeostasis. As expected, under both treatments, SD increased the amount of NREM-SWA compared to baseline sleep. Moreover, recovery sleep facilitated the dissipation of NREM-SWA to baseline levels for both treatments. Additionally, the spectral power of the lower frequency bands in the delta range (1-2.5 hertz) were significantly increased following SD for both treatments. However, analysis of the percent change for NREM-SWA (both amount and spectral power distribution) from baseline sleep revealed that E2 treatment compared to control resulted in a greater magnitude of change following sleep deprivation. Quantification of the AUC for the spectral power suggested that E2 induced an almost 2-fold increase in delta power in SD recovery sleep. The magnitude of E2 effects was a function of decreased levels of NREM-SWA in AL sleep suggesting that E2 significantly expands the dynamic range for the sleep homeostat. Increased wake at the expense of NREM sleep during the SD recovery following E2 treatment supports the assertion that E2 may expand the sleep homeostat set-points for sleep-need. Support for E2 regulation of homeostatic processes is clearly demonstrated in thermoregulatory process underlying vasomotor flushes in women. Loss of E2 significantly narrows thermal neutral zone which is the range of core body temperatures where body temperatures are maintained with minimal energy expenditure (i.e. sweating or shivering).^76–78^ Thus, in menopausal women where ovarian estrogens are severely depleted extremely small elevations in core body temperatures elicits a vasomotor flushes. Perhaps even more intriguing to the current study is that rodent studies have identified the temperature sensing neurons in the MnPO as site of E2 regulation.^79–83^

### Estradiol and Adenosine

The MnPO is a putative site for the sleep inducing actions of adenosine as intracerebroventricular injections of an A_2A_R agonist increase NREM sleep duration, sleep propensity and NREM-SWA, and activation of sleep-active GABAergic neurons.^61, 84^ Nevertheless, the current finding that E2 markedly increased MnPO extracellular adenosine during AL and SD recovery sleep in the absence of the hallmarks of increased sleep pressure presents a curious paradox about adenosine action in the presence of E2. Interestingly, other groups have reported increased wake behavior correlated with marked increases in POA adenosine.^84, 85^ When levels of extracellular adenosine were experimentally increased by either infusion of nitrobenzylthioinosine (an adenosine transport inhibitor) or infusion of high concentrations of adenosine, NREM sleep is reduced, and wake increased in male rats. Taken together with our current results, these findings raise a critical question concerning adenosinergic receptor signaling in the presence of elevated adenosine levels; specifically whether changes in adenosinergic signaling in the MnPO underlie E2-induced sleep suppression. Of note, another recent study has reported sex and estrous cycle dependent differences in adenosine release in the hippocampus, basal lateral amygdala and prefrontal cortex also suggesting that E2 may increase release.^86^

### Estradiol and Adenosine A_2A_ Receptors

The current findings suggest that E2 blocked the sleep-inducing effects of the A_2A_ receptor agonist, CGS, when the agonist was locally infused in the MnPO. The central somnogenic actions of adenosine occur via the A_1_ and A_2A_ receptors located in brain nuclei associated with sleep-wake behaviors. Sleep induction via A_1_R occurs through the inhibition of several wake-promoting areas including the cholinergic arousal system in the brainstem^87^ and basal forebrain^88^ the orexinergic system in the lateral hypothalamus^89^ and the histaminergic system in the tuberomammillary nucleus in the posterior hypothalamus.^90^ In contrast, more recent evidence suggests that A_2A_ receptors also play a significant role in sleep induction by exciting the GABAergic sleep-active neurons residing in the ventrolateral preoptic area (VLPO)^91^ and the median preoptic nucleus (MnPO).^49, 61^ Classic pharmacological experiments in male rats demonstrate that infusion of a highly selective A_1_R agonist, N^6^-cyclo-pentyladenosine (CPA) into the basal forebrain increases NREM and REM sleep. Subsequent work focusing on the POA sleep active nuclei suggest that activation of A_2A_ receptor with CGS stimulates sleep-active neurons in the VLPO^94^ and MnPO^61^ and increases GABA release in key arousal centers like the TMN.^95^ Of note in the current study is the possibility that the local CGS infusion may have had additional off-target (i.e. outside of the MnPO) effects that contributed to the increase in NREM-sleep. In a set of prior experiments, CGS was infused intraventricularly into adult OVX female rats in the absence and presence of exogenously administered sub threshold doses of E2 and results were identical to those presented here (Data not shown). Nevertheless, given our findings that the MnPO is a key site of E2 action, the current experiments further support its role in the estrogenic mechanisms governing sleep-wake behavior.

Interestingly, activation of POA A_1_ receptors by local infusion of CPA markedly increases wake at the expense of NREM sleep.^84^ Indeed, the observation that activation of POA A_1_ receptors induce wake may offer a possible insight into how E2 is increasing wake in the presence of increased adenosine; E2 may be shifting the adenosinergic balance of the excitatory A_2A_ tone that activates the GABAergic sleep-active neurons to an inhibitory A_1_ tone which inhibits the sleep-active neuronal population in the MnPO. The current findings suggest that E2 attenuated the signaling of the MnPO A_2A_ receptors when activated by CGS. However, the E2 was administered at subthreshold levels to prevent behavioral changes in sleep-wake. Thus, it remains unclear whether E2-induced increases in adenosine activate A_1_ receptors leading to an increase in wake. These experiments are currently on-going.

While the exact mechanism for how elevated levels of extracellular adenosine induce wake is not known, previous findings do suggest several possibilities including a shift in the excitatory/inhibitory balance^96^; direct post synaptic actions on sleep-neurons such as an uncoupling/downregulation of A_2A_ signaling^84^ or A_1_ mediated disinhibition of GABAergic inputs to sleep-active neurons. Our current findings strongly suggest that the E2-induced increases in extracellular MnPO adenosine may be the upstream-trigger of these potential subsequent actions on the signaling inputs that result in an inhibition of MnPO sleep neurons and increases in wake (Figure 8). Future work will seek to determine whether E2 requires a marked increase in extracellular MnPO adenosine to mediate its effects on sleep.

**Figure 8.**
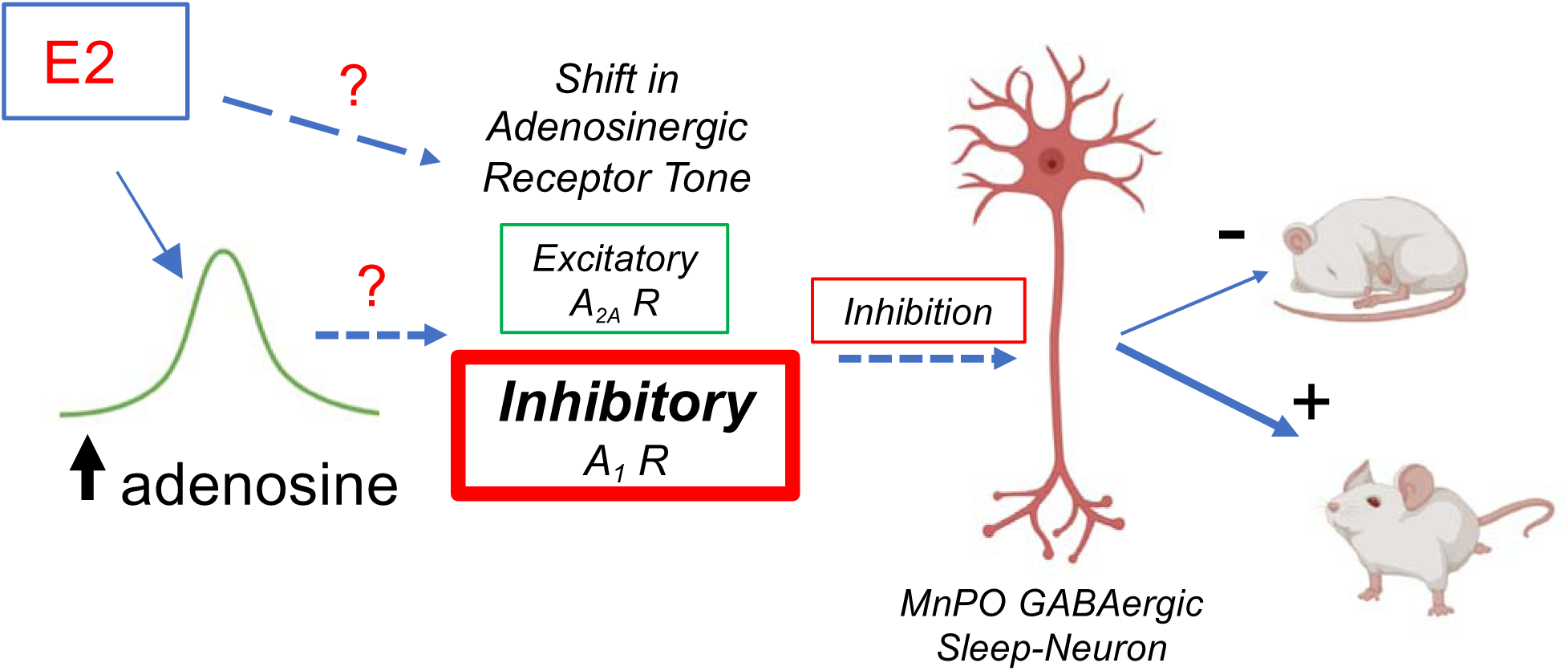
Proposed model of E2 action. The current findings demonstrate the E2 treatment increases extracellular levels of adenosine in the MnPO during both normal sleep condition and sleep deprivation. This increase in adenosine is associated with E2 induced decreases in sleep behavior or homeostatic pressure for sleep. Moreover, in the presence of E2 the ability to activate the A_2A_ receptor to induce sleep is attenuated. When taken together with evidence in the literature that, in the preoptic area, induced increases in extracellular adenosine or activation of the adenosine A_1_ receptor also increase wake and decrease NREM, our data may suggest that E2 is acting to shift the balance of adenosinergic signaling to a more inhibitory tone that inhibits sleep active neurons leading to increases in wake and/or decreases in homeostatic sleep pressure. It is not clear if E2 is directly acting to shift this balance or if the increase in adenosine is the trigger.

### Conclusions

While sex steroids and biological sex have been implicated as risk factors for sleep disruptions and insomnia, the relationship between ovarian steroids and normal sleep continues to be under investigated and poorly understood. Moreover, the mechanisms mediating ovarian steroid control of sleep are unknown. The current findings begin to lay the foundation for understanding potential cellular mechanisms underlying estrogenic effects on vigilance states and possibly sleep homeostasis by demonstrating a novel interaction between E2 and adenosinergic signaling in a major sleep-active nucleus, the MnPO. Future work will continue to elucidate the significance of this E2- adenosine nexus on regulation of sleep. Understanding the role of estrogenic regulation of sleep mechanisms is a critical first step toward a better understanding of its role in sleep pathologies and ultimately identification of targets for improved interventions for treating sleep disturbances in women and men.

## Acknowledgements

This research was funded by NIH grant #1F30HL145901 (granted to PCS) and grant # 5R01HL129138 (granted to JAM).

Adenosine measurement was performed at University of Maryland School of Medicine Center for Innovative Biomedical Resources, Protein Analysis Core – Baltimore, Md. LC/MS was performed at the Molecular Characterization and Analysis Complex at the University of Maryland, Baltimore County, Arbutus, Md.

The scoring algorithm was developed with programming assistance from Dr. Michael Rempe, Department of Mathematics and Computer Science, Whitworth University, Spokane, Wash.

---

**Supplementary Figure S1:**
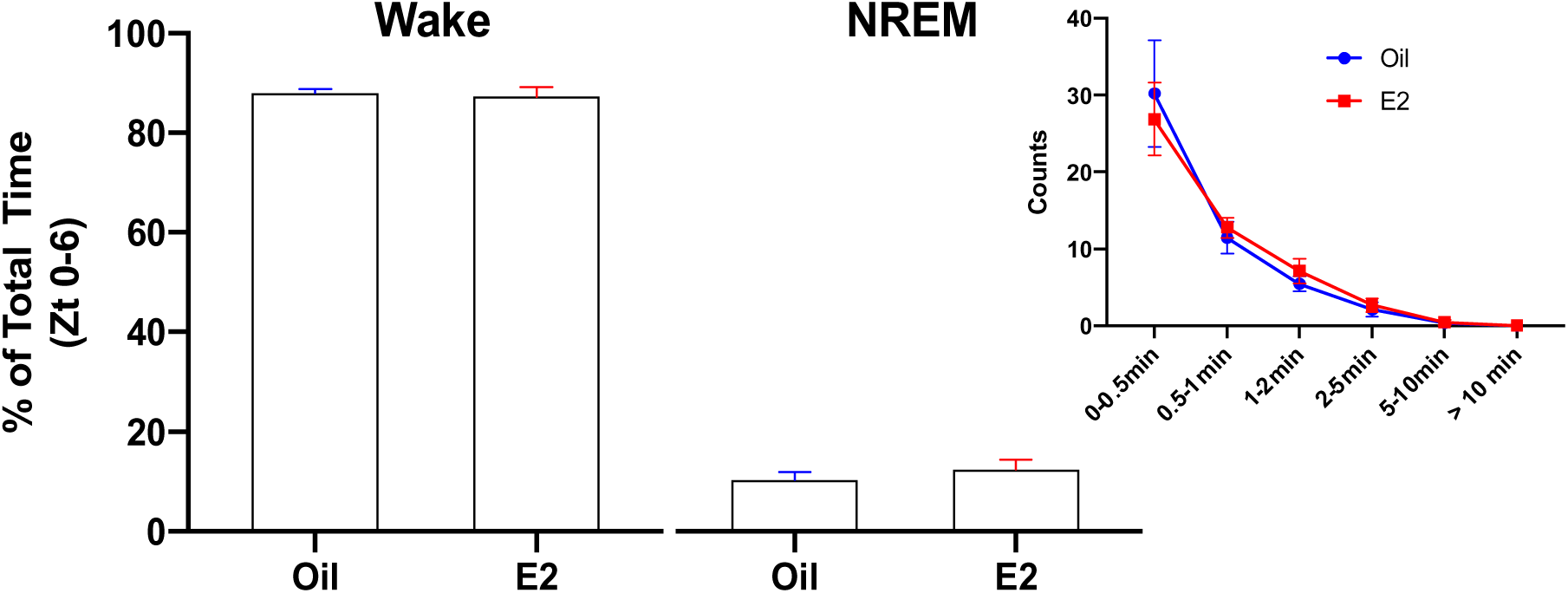
Sleep Deprivation Efficiency. Sleep deprivation was induced in the animal’s home cage between Zt 0-6. Analysis of the percent of total time spent in wake, NREM sleep and REM sleep over the 6-hour deprivation period was quantified for each animal following E2 or oil treatment from the associated EEG/EMG traces. The mean percent for each state was compared between the E2 and oil treatment via a paired t-test. The amount of accumulated Wake, NREM sleep an REM sleep was similar for both Oil and E2 treatment days (two-tailed paired t-test: Wake: p=0.67, t_(8)_=0.444; NREM: p=0.34, t_(8)_=1.026; REM: p=0.26, t_(8)_=1.23). The majority of accumulated NREM sleep occurred in bouts that were less than 1 minute and most often less than 30 seconds (inset). There was less than 0.1% of accumulated REM in either group (Data not shown). Data are the mean±SEM.

**Supplementary Figure S2:**
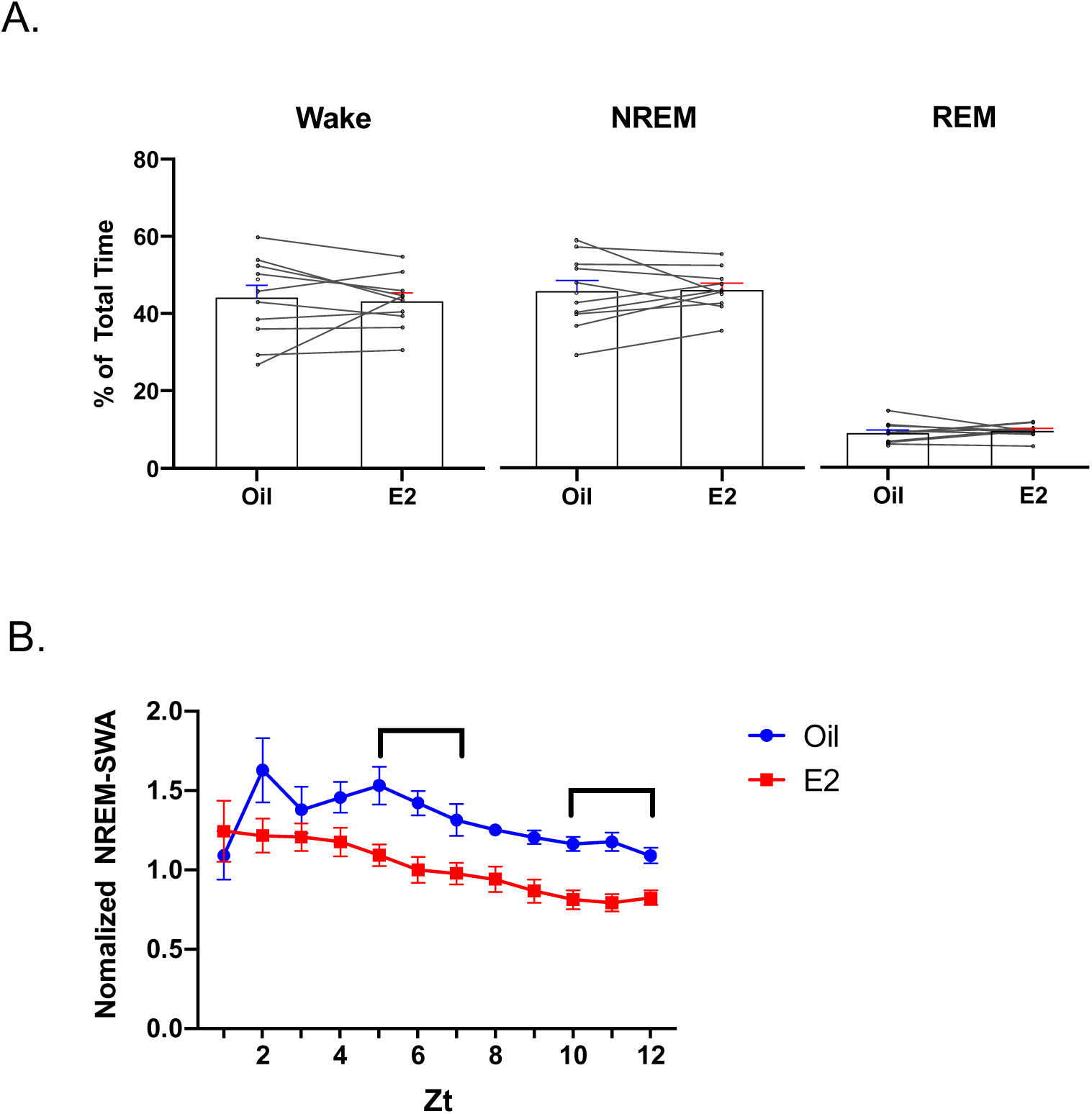
AL sleep 12-hour light phase totals for sleep-wake durations and NREM-SWA. (A) *E2 did not significantly affect the time spent in any state.* When the entire Day 3 light phase (Zt 1-12) was analyzed for the time spent in each vigilant state, E2 treatment did not significantly affect the percent of total time spent in any one state compared to oil. (B) *E2 reduced NREM-SWA across the light phase.* On Day 3, SWA during NREM sleep was analyzed in animals sleeping ad libitum for Zt 0-12. E2 treated animals showed significantly lower NREM-SWA across the entire light phase. (Repeated Measure two-way ANOVA; main effect of E2; F_(1, 84)_= 68.82, p<0.0001). A post hoc multiple comparison revealed significant differences between E2 and oil treatment at several timepoints throughout the light phase (Sidak’s multiple comparison test; p<0.05 Zt 5, Zt 6, Zt 7, Zt 10, Zt 11, Zt 12). Data are the mean±SEM.

**Supplementary Figure S3:**
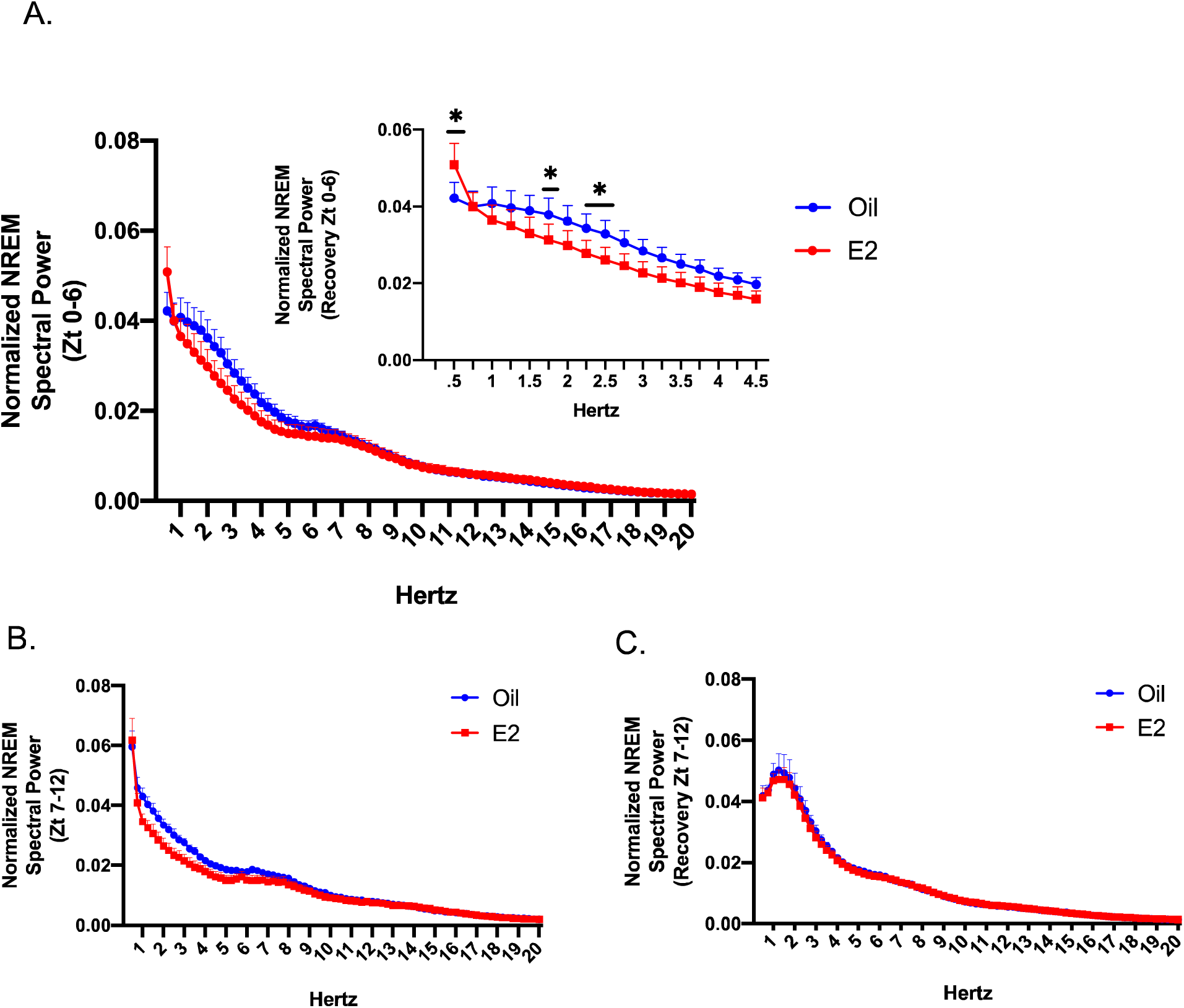
Distribution of spectral power following E2 and oil treatments in AL and SD recovery sleep. (A) In AL sleep, analysis of the spectral power distribution for NREM sleep during the first half of the light phase (Zt 0-6) revealed that E2-induced reductions in spectral power compared to oil were limited to the lower frequency delta range. A repeated measure mixed-effect analysis found a main effect of E2 (F_1,632_ = 14.14, p=0.0002) a main effect of frequency (F_78, 711_= 42.35, p<0.0001) and an interaction between the two (F_78,632_=2.034, p<0.0001). A post hoc multiple comparison test revealed significant differences between E2 and oil treatment at frequencies 2.5 Hz and below which is represented in the inset (Sidak’s multiple comparison test; *p<0.05 0.5Hz, 1.25Hz, 2.25Hz and 2.5Hz). (B,C) Expanded representation of the spectral power distribution in NREM sleep for Zt 7-12 in AL sleep (B) and SD recovery sleep (C). The delta range frequencies are presented in Figure 3. In AL sleep, analysis of the spectral power distribution for NREM sleep during the first half of the light phase (Zt 0-6) revealed that E2-induced reductions in spectral power compared to oil were limited to the lower frequency delta range. A repeated measure mixed-effect analysis found a main effect of E2 (F_1,632_ = 14.14, p=0.0002) a main effect of frequency (F_78, 711_= 42.35, p<0.0001) and an interaction between the two (F_78,632_=2.034, p<0.0001). A post hoc multiple comparison test revealed significant differences between E2 and oil treatment at frequencies 2.5 Hz and below which is represented in the inset (Sidak’s multiple comparison test; *p<0.05 0.5Hz, 1.25Hz, 2.25Hz and 2.5Hz). Data are the mean±SEM.

**Supplementary Figure S4:**
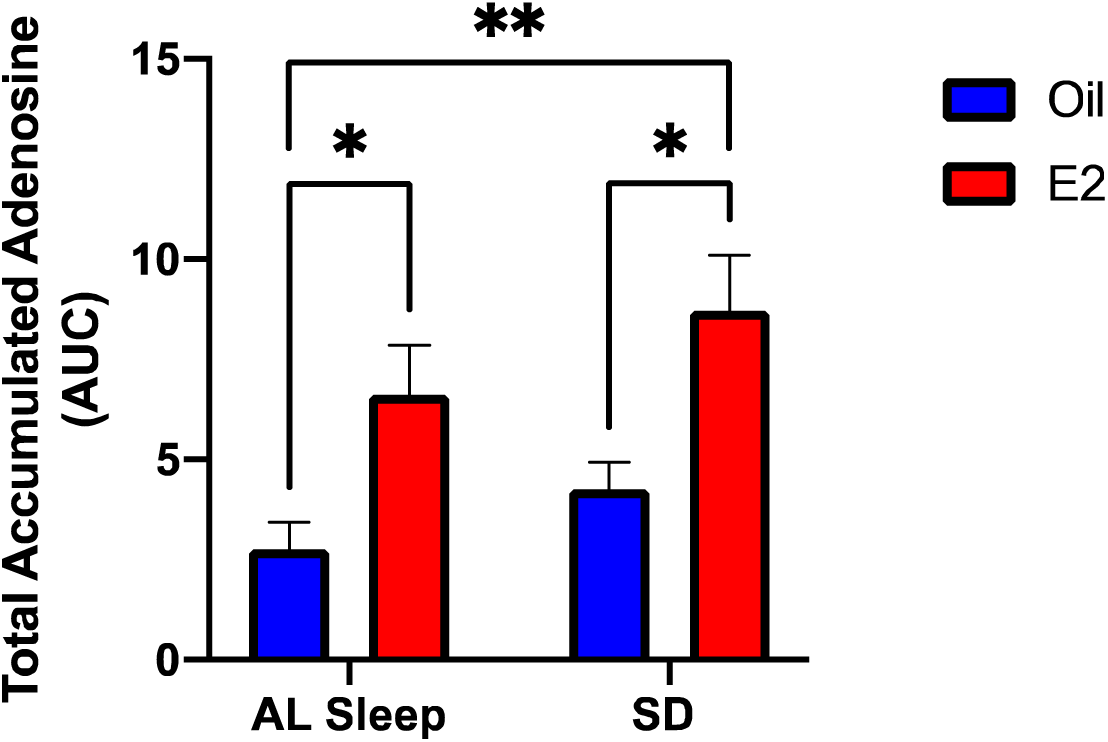
Area under the curve comparison of accumulated extracellular adenosine levels between the AL and SD sleep experiments. Comparing the AUC calculations across the four treatment groups demonstrated that SD did increase total adenosine content in both the oil and E2 treated groups compared to AL sleep; however, the increase did not reach statistical significance (Two way ANOVA; main effect of E2; F_1,29_= 15.96, p=0.0004; no main effect of sleep condition F_1,29_ = 2.961, p=0.0960). This could be due to the experimental variability between the AL and SD cohorts which were run as separate experiments. Post hoc analysis revealed that significant differences between E2 vs oil in AL sleep and SD (Sidak’s multiple comparison test p=0.0381 and p=0.0402, respectively) as well as E2 in SD vs Oil in AL (p=0.0017). Data are the mean±SEM.

**Supplementary Table 1:**
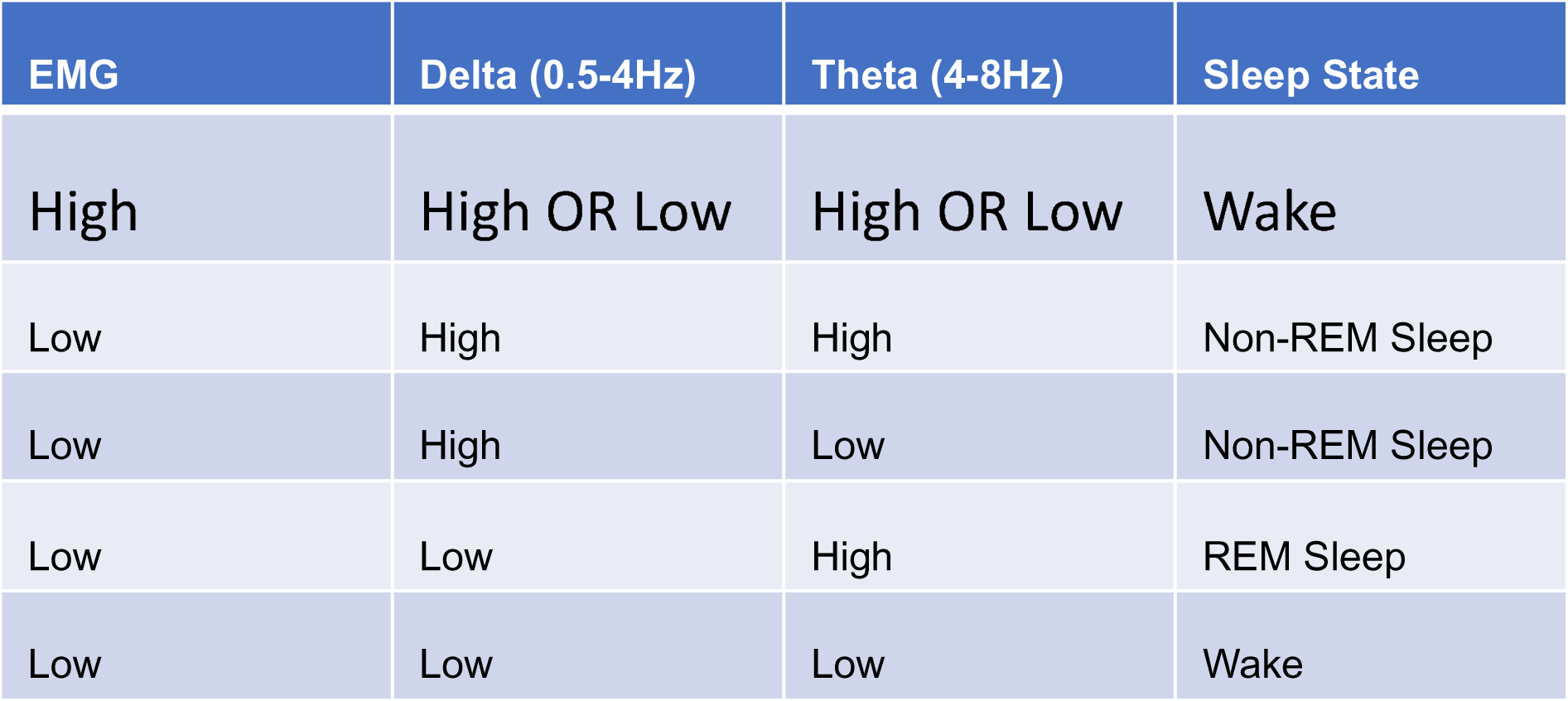
MATLAB Automated Vigilant State Scoring. Wake, NREM sleep and REM sleep were assigned to each 10 second epoch based on threshold calls of muscle tone (EMG) and Delta/Theta power. Determinations of the low or high thresholds were made relative to each animal’s median value for the given parameter.

**Supplementary Table 2:**
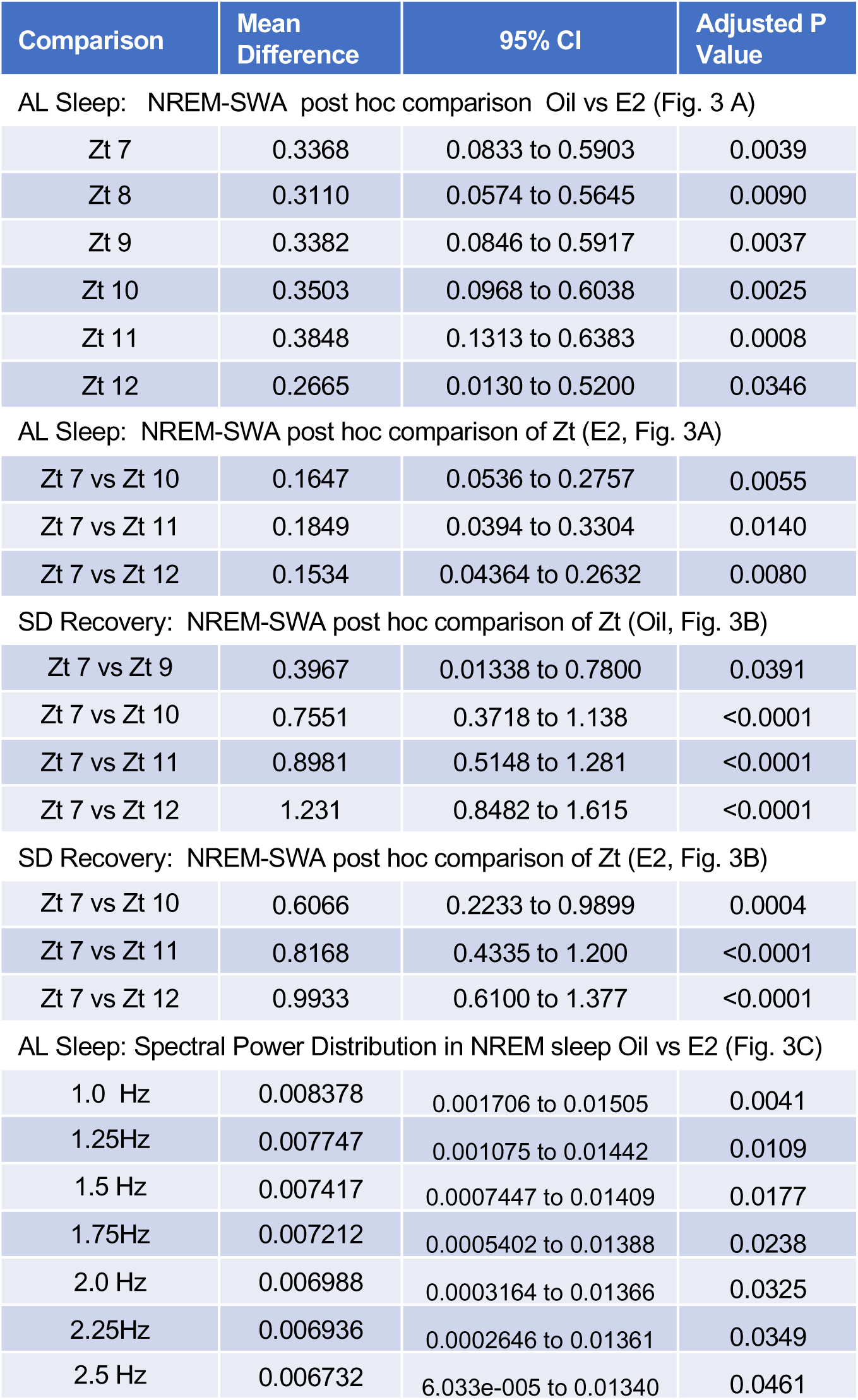
Sidak’s multiple comparison test Adjusted P values and confidence intervals for Figure 3. List of the mean difference, confidence intervals (CI) and adjusted P values for the comparisons in Figure 3.

| Comparison | Mean Difference | 95% CI | Adjusted P Value |
| --- | --- | --- | --- |
| AL Sleep: NREM-SWA post hoc comparison Oil vs E2 (Fig. 3 A) |  |  |  |
| Zt 7 | 0.3368 | 0.0833 to 0.5903 | 0.0039 |
| Zt 8 | 0.3110 | 0.0574 to 0.5645 | 0.0090 |
| Zt 9 | 0.3382 | 0.0846 to 0.5917 | 0.0037 |
| Zt 10 | 0.3503 | 0.0968 to 0.6038 | 0.0025 |
| Zt 11 | 0.3848 | 0.1313 to 0.6383 | 0.0008 |
| Zt 12 | 0.2665 | 0.0130 to 0.5200 | 0.0346 |
| AL Sleep: NREM-SWA post hoc comparison of Zt (E2, Fig. 3A) |  |  |  |
| Zt 7 vs Zt 10 | 0.1647 | 0.0536 to 0.2757 | 0.0055 |
| Zt 7 vs Zt 11 | 0.1849 | 0.0394 to 0.3304 | 0.0140 |
| Zt 7 vs Zt 12 | 0.1534 | 0.04364 to 0.2632 | 0.0080 |
| SD Recovery: NREM-SWA post hoc comparison of Zt (Oil, Fig. 3B) |  |  |  |
| Zt 7 vs Zt 9 | 0.3967 | 0.01338 to 0.7800 | 0.0391 |
| Zt 7 vs Zt 10 | 0.7551 | 0.3718 to 1.138 | <0.0001 |
| Zt 7 vs Zt 11 | 0.8981 | 0.5148 to 1.281 | <0.0001 |
| Zt 7 vs Zt 12 | 1.231 | 0.8482 to 1.615 | <0.0001 |
| SD Recovery: NREM-SWA post hoc comparison of Zt (E2, Fig. 3B) |  |  |  |
| Zt 7 vs Zt 10 | 0.6066 | 0.2233 to 0.9899 | 0.0004 |
| Zt 7 vs Zt 11 | 0.8168 | 0.4335 to 1.200 | <0.0001 |
| Zt 7 vs Zt 12 | 0.9933 | 0.6100 to 1.377 | <0.0001 |

|  |  |  |  |
| --- | --- | --- | --- |
| 1.0 Hz | 0.008378 | 0.001706 to 0.01505 | 0.0041 |
| 1.25Hz | 0.007747 | 0.001075 to 0.01442 | 0.0109 |
| 1.5 Hz | 0.007417 | 0.0007447 to 0.01409 | 0.0177 |
| 1.75Hz | 0.007212 | 0.0005402 to 0.01388 | 0.0238 |
| 2.0 Hz | 0.006988 | 0.0003164 to 0.01366 | 0.0325 |
| 2.25Hz | 0.006936 | 0.0002646 to 0.01361 | 0.0349 |
| 2.5 Hz | 0.006732 | 6.033e-005 to 0.01340 | 0.0461 |

## References

1. Ford ES, Cunningham TJ, Croft JB. Trends in Self-Reported Sleep Duration among US Adults from 1985 to 2012. Sleep. 2015; 38 (5): 829–832.

2. Ford ES, Cunningham TJ, Giles WH, Croft JB. Trends in insomnia and excessive daytime sleepiness among U.S. adults from 2002 to 2012. Sleep Med. 2015; 16 (3): 372–378.

3. Watson NF, Badr MS, Belenky G, et al. Joint Consensus Statement of the American Academy of Sleep Medicine and Sleep Research Society on the Recommended Amount of Sleep for a Healthy Adult: Methodology and Discussion. J Clin Sleep Med. 2015; 11 (8): 931–952.

4. Wickwire EM, Shaya FT, Scharf SM. Health economics of insomnia treatments: The return on investment for a good night’s sleep. Sleep Med Rev. 2016; 30: 72–82.

5. Ohayon MM. Prevalence of DSM-IV diagnostic criteria of insomnia: distinguishing insomnia related to mental disorders from sleep disorders. Journal of psychiatric research. 1997; 31 (3): 333–346.

6. Hu X-Z, Lipsky RH, Zhu G, et al. Serotonin transporter promoter gain-of-function genotypes are linked to obsessive-compulsive disorder. Am J Hum Genet. 2006; 78 (5): 815–826.

7. Kravitz HM, Joffe H. Sleep during the perimenopause: a SWAN story. Obstet Gynecol Clin North Am. 2011; 38 (3): 567–586.

8. Johnson EO, Roth T, Schultz L, Breslau N. Epidemiology of DSM-IV insomnia in adolescence: lifetime prevalence, chronicity, and an emergent gender difference. Pediatrics. 2006; 117 (2): e247–256.

9. Zhang J, Chan NY, Lam SP, et al. Emergence of Sex Differences in Insomnia Symptoms in Adolescents: A Large-Scale School-Based Study. Sleep. 2016; 39 (8): 1563–1570.

10. Mong JA, Cusmano DM. Sex differences in sleep: impact of biological sex and sex steroids. Philosophical Transactions of the Royal Society B: Biological Sciences. 2016; 371 (1688): 20150110.

11. Parry BL, Mendelson WB, Duncan WC, Sack DA, Wehr TA. Longitudinal sleep EEG, temperature, and activity measurements across the menstrual cycle in patients with premenstrual depression and in age-matched controls. Psychiatry research. 1989; 30 (3): 285–303.

12. Dzaja A, Arber S, Hislop J, et al. Women’s sleep in health and disease. Journal of psychiatric research. 2005; 39 (1): 55–76.

13. Joffe H, Massler A, Sharkey KM. Evaluation and management of sleep disturbance during the menopause transition. Semin Reprod Med. 2010; 28 (5): 404–421.

14. Shechter A, Boivin DB. Sleep, Hormones, and Circadian Rhythms throughout the Menstrual Cycle in Healthy Women and Women with Premenstrual Dysphoric Disorder. International journal of endocrinology. 2010; 2010: 259345.

15. Shechter A, Varin F, Boivin DB. Circadian variation of sleep during the follicular and luteal phases of the menstrual cycle. Sleep. 2010; 33 (5): 647–656.

16. Redline S, Kump K, Tishler PV, Browner I, Ferrette V. Gender differences in sleep disordered breathing in a community-based sample. Am J Respir Crit Care Med. 1994; 149 (3 Pt 1): 722–726.

17. Mendez M, Radtke RA. Interactions between sleep and epilepsy. Journal of clinical neurophysiology : official publication of the American Electroencephalographic Society. 2001; 18 (2): 106–127.

18. Vock J, Achermann P, Bischof M, et al. Evolution of sleep and sleep EEG after hemispheric stroke. J Sleep Res. 2002; 11 (4): 331–338.

19. Petit D, Gagnon JF, Fantini ML, Ferini-Strambi L, Montplaisir J. Sleep and quantitative EEG in neurodegenerative disorders. J Psychosom Res. 2004; 56 (5): 487–496.

20. Somers VK, White DP, Amin R, et al. Sleep apnea and cardiovascular disease: an American Heart Association/american College Of Cardiology Foundation Scientific Statement from the American Heart Association Council for High Blood Pressure Research Professional Education Committee, Council on Clinical Cardiology, Stroke Council, and Council On Cardiovascular Nursing. In collaboration with the National Heart, Lung, and Blood Institute National Center on Sleep Disorders Research (National Institutes of Health). Circulation. 2008; 118 (10): 1080–1111.

21. Pasic Z, Smajlovic D, Dostovic Z, Kojic B, Selmanovic S. Incidence and types of sleep disorders in patients with stroke. Medicinski arhiv. 2011; 65 (4): 225–227.

22. Ju YE, Lucey BP, Holtzman DM. Sleep and Alzheimer disease pathology--a bidirectional relationship. Nat Rev Neurol. 2014; 10 (2): 115–119.

23. Musiek ES, Holtzman DM. Mechanisms linking circadian clocks, sleep, and neurodegeneration. Science. 2016; 354 (6315): 1004–1008.

24. Ayas NT, White DP, Manson JE, et al. A prospective study of sleep duration and coronary heart disease in women. Arch Intern Med. 2003; 163 (2): 205–209.

25. Meier-Ewert HK, Ridker PM, Rifai N, et al. Effect of sleep loss on C-reactive protein, an inflammatory marker of cardiovascular risk. J Am Coll Cardiol. 2004; 43 (4): 678–683.

26. Patel SR, Ayas NT, Malhotra MR, et al. A prospective study of sleep duration and mortality risk in women. Sleep. 2004; 27 (3): 440–444.

27. Gangwisch JE, Heymsfield SB, Boden-Albala B, et al. Short sleep duration as a risk factor for hypertension: analyses of the first National Health and Nutrition Examination Survey. Hypertension. 2006; 47 (5): 833–839.

28. Cappuccio FP, Stranges S, Kandala NB, et al. Gender-specific associations of short sleep duration with prevalent and incident hypertension: the Whitehall II Study. Hypertension. 2007; 50 (4): 693–700.

29. Krystal AD. Depression and insomnia in women. Clin Cornerstone. 2004; 6 Suppl 1B: S19–28.

30. Lyytikainen P, Rahkonen O, Lahelma E, Lallukka T. Association of sleep duration with weight and weight gain: a prospective follow-up study. Journal of sleep research. 2011; 20 (2): 298–302.

31. Ferrie JE, Shipley MJ, Cappuccio FP, et al. A prospective study of change in sleep duration: associations with mortality in the Whitehall II cohort. Sleep. 2007; 30 (12): 1659–1666.

32. Miller MA, Kandala NB, Kivimaki M, et al. Gender differences in the cross-sectional relationships between sleep duration and markers of inflammation: Whitehall II study. Sleep. 2009; 32 (7): 857–864.

33. Kronholm E, Laatikainen T, Peltonen M, Sippola R, Partonen T. Self-reported sleep duration, all-cause mortality, cardiovascular mortality and morbidity in Finland. Sleep medicine. 2011; 12 (3): 215–221.

34. Merklinger-Gruchala A, Ellison PT, Lipson SF, Thune I, Jasienska G. Low estradiol levels in women of reproductive age having low sleep variation. European journal of cancer prevention : the official journal of the European Cancer Prevention Organisation. 2008; 17 (5): 467–472.

35. Baker FC, O’Brien LM, Armitage R. Sex differences and menstrual-related changes in sleep and circadian rhythms. In: Kryger M, Roth T, Dement WC, eds. Principles and Practice of Sleep Medicine (5th Ed). St. Louis, Missouri: Elsevier Saunders.; 2010: 1562–1571.

36. Mong JA, Baker FC, Mahoney MM, et al. Sleep, rhythms, and the endocrine brain: influence of sex and gonadal hormones. J Neurosci. 2011; 31 (45): 16107–16116.

37. Shechter A, Lesperance P, Ng Ying Kin NM, Boivin DB. Nocturnal polysomnographic sleep across the menstrual cycle in premenstrual dysphoric disorder. Sleep Med. 2012; 13 (8): 1071–1078.

38. Sharkey KM, Crawford SL, Kim S, Joffe H. Objective sleep interruption and reproductive hormone dynamics in the menstrual cycle. Sleep Med. 2014; 15 (6): 688–693.

39. Schwierin B, Borbely AA, Tobler I. Prolonged effects of 24-h total sleep deprivation on sleep and sleep EEG in the rat. Neuroscience Letters. 1999; 261 (1-2): 61–64.

40. Hadjimarkou MM, Benham R, Schwarz JM, Holder MK, Mong JA. Estradiol suppresses rapid eye movement sleep and activation of sleep-active neurons in the ventrolateral preoptic area. Eur J Neurosci. 2008; 27 (7): 1780–1792.

41. Swift KM, Keus K, Echeverria CG, et al. Sex differences within sleep in gonadally intact rats. Sleep. 2020; 43 (5).

42. Cusmano DM, Hadjimarkou MM, Mong JA. Gonadal steroid modulation of sleep and wakefulness in male and female rats is sexually differentiated and neonatally organized by steroid exposure. Endocrinology. 2014; 155 (1): 204–214.

43. Mong JA, Devidze N, Frail DE, et al. Estradiol differentially regulates lipocalin-type prostaglandin D synthase transcript levels in the rodent brain: Evidence from high-density oligonucleotide arrays and in situ hybridization. PNAS. 2003; 100 (1): 318–323.

44. Mong JA, Devidze N, Goodwillie A, Pfaff DW. Reduction of lipocalin-type prostaglandin D synthase in the preoptic area of female mice mimics estradiol effects on arousal and sex behavior. Proceedings of the National Academy of Sciences of the United States of America. 2003; 100 (25): 15206–15211.

45. Smith PC, Phillips DJ, Viechweg SS, Schwartz MD, Mong JA. Estradiol Action at the Median Preoptic Nucleus is Necessary and Sufficient for Sleep Suppression in Female Rats. bioRxiv. 2020: 2020.2007.2029.223669.

46. Lu J, Greco MA, Shiromani P, Saper CB. Effect of Lesions of the Ventrolateral Preoptic Nucleus on NREM and REM Sleep. J Neurosci. 2000; 20 (10): 3830–3842.

47. Gvilia I, Angara C, McGinty D, Szymusiak R. Different neuronal populations of the rat median preoptic nucleus express c-fos during sleep and in response to hypertonic saline or angiotensin-II. The Journal of physiology. 2005; 569 (Pt 2): 587–599.

48. Gvilia I, Xu F, McGinty D, Szymusiak R. Homeostatic regulation of sleep: a role for preoptic area neurons. The Journal of neuroscience : the official journal of the Society for Neuroscience. 2006; 26 (37): 9426–9433.

49. Alam MA, Kumar S, McGinty D, Alam MN, Szymusiak R. Neuronal activity in the preoptic hypothalamus during sleep deprivation and recovery sleep. J Neurophysiol. 2014; 111 (2): 287–299.

50. Borbely AA, Baumann F, Brandeis D, Strauch I, Lehmann D. Sleep deprivation: effect on sleep stages and EEG power density in man. Electroencephalogr Clin Neurophysiol. 1981; 51 (5): 483–495.

51. Franken P, Dijk DJ, Tobler I, Borbely AA. Sleep deprivation in rats: effects on EEG power spectra, vigilance states, and cortical temperature. Am J Physiol. 1991; 261 (1 Pt 2): R198–208.

52. Franken P, Tobler I, Borbely AA. Sleep homeostasis in the rat: simulation of the time course of EEG slow-wave activity. Neurosci Lett. 1991; 130 (2): 141–144.

53. Borbely AA. Sleep mechanisms. Sleep and Biological Rhythms. 2004; 2: S67– S68.

54. Borbely AA, Tobler I. Manifestations and functional implications of sleep homeostasis. Handb Clin Neurol. 2011; 98: 205–213.

55. Deboer T. Behavioral and electrophysiological correlates of sleep and sleep homeostasis. Curr Top Behav Neurosci. 2015; 25: 1–24.

56. Lazarus M, Chen JF, Huang ZL, Urade Y, Fredholm BB. Adenosine and Sleep. Handb Exp Pharmacol. 2019; 253: 359–381.

57. Lazarus M, Oishi Y, Bjorness TE, Greene RW. Gating and the Need for Sleep: Dissociable Effects of Adenosine A1 and A2A Receptors. Front Neurosci. 2019; 13: 740.

58. Porkka-Heiskanen T, Strecker RE, Thakkar M, Bjorkum AA, Greene RW, McCarley RW. Adenosine: a mediator of the sleep-inducing effects of prolonged wakefulness. Science. 1997; 276 (5316): 1265–1268.

59. Porkka-Heiskanen T, Strecker RE, McCarley RW. Brain site-specificity of extracellular adenosine concentration changes during sleep deprivation and spontaneous sleep: an in vivo microdialysis study. Neuroscience. 2000; 99 (3): 507–517.

60. Strecker RE, Morairty S, Thakkar MM, et al. Adenosinergic modulation of basal forebrain and preoptic/anterior hypothalamic neuronal activity in the control of behavioral state. Behav Brain Res. 2000; 115 (2): 183–204.

61. Kumar S, Rai S, Hsieh KC, McGinty D, Alam MN, Szymusiak R. Adenosine A(2A) receptors regulate the activity of sleep regulatory GABAergic neurons in the preoptic hypothalamus. Am J Physiol Regul Integr Comp Physiol. 2013; 305 (1): R31–41.

62. Schwierin B, Borbely AA, Tobler I. Sleep homeostasis in the female rat during the estrous cycle. Brain Res. 1998; 811 (1-2): 96–104.

63. Schwartz MD, Mong JA. Estradiol modulates recovery of REM sleep in a time-of-day-dependent manner. Am J Physiol Regul Integr Comp Physiol. 2013; 305 (3): R271–280.

64. Oriowo MA, Landgren BM, Stenstrom B, Diczfalusy E. A comparison of the pharmacokinetic properties of three estradiol esters. Contraception. 1980; 21 (4): 415–424.

65. Lund-Pero M, Jeppson B, Arneklo-Nobin B, Sjogren HO, Holmgren K, Pero RW. Non-specific steroidal esterase activity and distribution in human and other mammalian tissues. Clin Chim Acta. 1994; 224 (1): 9–20.

66. Paxinos G, Watson C. The rat brain in stereotaxic coordinates. Amsterdam; Boston: Elsevier Academic Press; 2005.

67. Schwartz MD, Mong JA. Estradiol suppresses recovery of REM sleep following sleep deprivation in ovariectomized female rats. Physiology & Behavior. 2011; 104 (5): 962–971.

68. Latini S, Pedata F. Adenosine in the central nervous system: release mechanisms and extracellular concentrations. J Neurochem. 2001; 79 (3): 463–484.

69. Blanco-Centurion C, Xu M, Murillo-Rodriguez E, et al. Adenosine and sleep homeostasis in the Basal forebrain. J Neurosci. 2006; 26 (31): 8092–8100.

70. Donlea JM, Alam MN, Szymusiak R. Neuronal substrates of sleep homeostasis; lessons from flies, rats and mice. Curr Opin Neurobiol. 2017; 44: 228–235.

71. Paul KN, Laposky AD, Turek FW. Reproductive hormone replacement alters sleep in mice. Neuroscience letters. 2009; 463 (3): 239–243.

72. Deurveilher S, Rusak B, Semba K. Estradiol and progesterone modulate spontaneous sleep patterns and recovery from sleep deprivation in ovariectomized rats. Sleep. 2009; 32 (7): 865–877.

73. Deurveilher S, Rusak B, Semba K. Female reproductive hormones alter sleep architecture in ovariectomized rats. Sleep. 2011; 34 (4): 519–530.

74. Holder MK, Hadjimarkou MM, Zup SL, et al. Methamphetamine facilitates female sexual behavior and enhances neuronal activation in the medial amygdala and ventromedial nucleus of the hypothalamus. Psychoneuroendocrinology. 2010; 35 (2): 197–208.

75. Holder MK, Mong JA. Methamphetamine enhances paced mating behaviors and neuroplasticity in the medial amygdala of female rats. Hormones & Behavior. 2010.

76. Freedman RR, Krell W. Reduced thermoregulatory null zone in postmenopausal women with hot flashes. Am J Obstet Gynecol. 1999; 181 (1): 66–70.

77. Freedman RR. Menopausal hot flashes: mechanisms, endocrinology, treatment. J Steroid Biochem Mol Biol. 2014; 142: 115–120.

78. Freedman RR. Postmenopausal physiological changes. Curr Top Behav Neurosci. 2014; 21: 245–256.

79. Dacks PA, Krajewski SJ, Rance NE. Ambient temperature and 17beta-estradiol modify Fos immunoreactivity in the median preoptic nucleus, a putative regulator of skin vasomotion. Endocrinology. 2011; 152 (7): 2750–2759.

80. Dacks PA, Krajewski SJ, Rance NE. Activation of neurokinin 3 receptors in the median preoptic nucleus decreases core temperature in the rat. Endocrinology. 2011; 152 (12): 4894–4905.

81. Mittelman-Smith MA, Krajewski-Hall SJ, McMullen NT, Rance NE. Neurokinin 3 Receptor-Expressing Neurons in the Median Preoptic Nucleus Modulate Heat-Dissipation Effectors in the Female Rat. Endocrinology. 2015; 156 (7): 2552–2562.

82. Krajewski-Hall SJ, Blackmore EM, McMinn JR, Rance NE. Estradiol alters body temperature regulation in the female mouse. Temperature (Austin). 2018; 5 (1): 56–69.

83. Krajewski-Hall SJ, Miranda Dos Santos F, McMullen NT, Blackmore EM, Rance NE. Glutamatergic Neurokinin 3 Receptor Neurons in the Median Preoptic Nucleus Modulate Heat-Defense Pathways in Female Mice. Endocrinology. 2019; 160 (4): 803–816.

84. Methippara MM, Kumar S, Alam MN, Szymusiak R, McGinty D. Effects on sleep of microdialysis of adenosine A1 and A2a receptor analogs into the lateral preoptic area of rats. Am J Physiol Regul Integr Comp Physiol. 2005; 289 (6): R1715–1723.

85. Zhang J, Yin D, Wu F, et al. Microinjection of adenosine into the hypothalamic ventrolateral preoptic area enhances wakefulness via the A1 receptor in rats. Neurochem Res. 2013; 38 (8): 1616–1623.

86. Borgus JR, Puthongkham P, Venton BJ. Complex Sex and Estrous Cycle Differences in Spontaneous Transient Adenosine. J Neurochem. 2020: e14981.

87. Rainnie DG, Grunze HC, McCarley RW, Greene RW. Adenosine inhibition of mesopontine cholinergic neurons: implications for EEG arousal. Science. 1994; 263 (5147): 689–692.

88. Alam MN, Szymusiak R, Gong H, King J, McGinty D. Adenosinergic modulation of rat basal forebrain neurons during sleep and waking: neuronal recording with microdialysis. J Physiol. 1999; 521 Pt 3: 679–690.

89. Martin LJ, Liu Z, Chen K, et al. Motor neuron degeneration in amyotrophic lateral sclerosis mutant superoxide dismutase-1 transgenic mice: mechanisms of mitochondriopathy and cell death. Journal of Comparative Neurology. 2007; 500 (1): 20–46.

90. Oishi Y, Huang ZL, Fredholm BB, Urade Y, Hayaishi O. Adenosine in the tuberomammillary nucleus inhibits the histaminergic system via A1 receptors and promotes non-rapid eye movement sleep. Proc Natl Acad Sci U S A. 2008; 105 (50): 19992–19997.

91. Scammell TE, Gerashchenko DY, Mochizuki T, et al. An adenosine A2a agonist increases sleep and induces Fos in ventrolateral preoptic neurons. Neuroscience. 2001; 107 (4): 653–663.

92. Benington JH, Kodali SK, Heller HC. Stimulation of A1 adenosine receptors mimics the electroencephalographic effects of sleep deprivation. Brain Res. 1995; 692 (1-2): 79–85.

93. Yang C, Franciosi S, Brown RE. Adenosine inhibits the excitatory synaptic inputs to Basal forebrain cholinergic, GABAergic, and parvalbumin neurons in mice. Frontiers in neurology. 2013; 4: 77.

94. Gallopin T, Luppi PH, Cauli B, et al. The endogenous somnogen adenosine excites a subset of sleep-promoting neurons via A2A receptors in the ventrolateral preoptic nucleus. Neuroscience. 2005; 134 (4): 1377–1390.

95. Hong CJ, Liu HC, Liu TY, Liao DL, Tsai SJ. Association studies of the adenosine A2a receptor (1976T > C) genetic polymorphism in Parkinson’s disease and schizophrenia. J Neural Transm (Vienna). 2005; 112 (11): 1503–1510.

96. Qi G, van Aerde K, Abel T, Feldmeyer D. Adenosine Differentially Modulates Synaptic Transmission of Excitatory and Inhibitory Microcircuits in Layer 4 of Rat Barrel Cortex. Cereb Cortex. 2017; 27 (9): 4411–4422.

